# A Modular Design for Synthetic Membraneless Organelles Enables Compositional and Functional Control

**DOI:** 10.1101/2023.10.03.560789

**Authors:** Mackenzie T. Walls, Ke Xu, Clifford P. Brangwynne, José L. Avalos

## Abstract

Living cells organize a wide variety of processes through compartmentalization into membraneless organelles, known as biomolecular condensates. Given their ubiquitous presence across a wide spectrum of different organisms and cell types, biomolecular condensates are increasingly considered to offer great potential for biotechnological applications. However, native condensates remain difficult to harness for engineering applications, both due to their intertwined mechanisms of assembly and compositional control, and potential disruptions to native cellular processes. Here, we demonstrate a modular framework for the formation of synthetic condensates designed to decouple cluster formation and protein recruitment. Synthetic condensates are built through constitutive oligomerization of intrinsically-disordered regions (IDRs), which drive the formation of condensates whose composition can be independently defined through fused interaction domains. The composition of the proteins driven to partition into the condensate can be quantitatively described using a binding equilibrium model, demonstrating predictive control of how component expression levels and interaction affinity determine the degree of protein recruitment. Finally, the engineered system is utilized to regulate protein interactions and metabolic flux by harnessing the system’s compositional tunability.

## Introduction

Subcellular compartmentalization is a pivotal feature of all known forms of life^1–3^. Within eukaryotic cells, canonical organelles such as mitochondria and chloroplasts use membranes to sequester specific cellular processes^4^. But organelles that lack enclosing lipid membranes have also received intense recent interest, and appear to be just as ubiquitous in eukaryotic cells^5,6^ and also prokaryotes^7,8^. These structures assemble through an array of self-interactions that lead to biomolecular phase separation or condensation^9–12^, similar to the demixing of immiscible liquids like oil and water, and are often referred to as biomolecular condensates^13,14^.

Biomolecular condensates are involved in a wide range of biological functions, each requiring unique compositions^15,16^. Broadly, condensates sequester specific biomolecules and exclude others to tune local concentrations, and therefore the resulting biomolecular interactions and kinetics^14,17^. Processes such as cellular signaling and enzymatic catalysis are sensitive to concentration amplification and are thus highly amenable to biological regulation through biomolecular condensation^18,19^. For example, purinosomes and G bodies enrich the enzymes catalyzing the metabolic cascades of purine biosynthesis^20^ and glycolysis^21,22^ to enhance pathway flux under states of high metabolic demand.

The broad range of native cellular functions that have been ascribed to condensates has motivated recent interest in harnessing these bodies for intracellular organelle engineering^6,23,24^. For instance, sequestration of proteins into a condensate can prevent their physiological localization^25^ and inhibit native interactions^26–28^. Alternatively, the recruitment of multiple proteins into a condensate can lead to a synergistic increase in biomolecular activity^28,29^. Inspired by purinosomes and G-bodies, recruitment of metabolic enzymes to synthetic condensates can enhance the output of their biosynthetic pathways^30–34^.

To disentangle and harness the principles behind functionalization and compositional control of native or engineered condensates, the enriched components are sometimes separated in two categories: scaffolds, which contribute to condensate assembly and architecture; and clients, which are cargo molecules that lack self-interactions necessary for phase separation, and simply interact with scaffolds to drive their recruitment^35^. However, native biomolecules often fail to fit squarely into one of these distinct categories. In particular, cargo recruitment is often intertwined with condensate assembly; an interplay which can regulate condensate formation and structure in subtle ways^36,37^, and potentially confound systematic manipulation of cargo recruitment. It is thus crucial to develop synthetic scaffold-client systems that exhibit defined interactions, uncoupling macromolecule condensation from cargo recruitment, and allowing for compositional control and tunable functionalization of condensates. Previous examples of synthetic membraneless organelles have pieced together protein chimeras with defined features to generate designer condensates^6,38,39^. For example, we previously introduced the Corelet system, whereby weakly interacting protein domains optogenetically multimerize on spherical protein assemblies of human ferritin, resulting in spatiotemporally tunable phase separation^40^. Others have added interaction domains to recruit various proteins to functionalize related synthetic organelles^26,27,33,34^. However, previous efforts to systematically engineer biomolecular condensates fall short of providing a quantitative framework for controlling the recruitment of various functional proteins into the condensates. Indeed, the physicochemical principles underlying the regulation of cargo composition in condensates remain largely unclear.

Here we develop a modular clustering and cargo recruitment system that we use to uncover and quantitatively map the design principles for condensate functionalization. We employ a range of peptide-interacting domains fused to condensate scaffolds to recruit protein cargo fused to the corresponding cognate peptide. With experimental data supported by theory, we develop a quantitative framework grounded in binding equilibrium whereby cargo concentration and affinity for the condensates act as tunable parameters to control the degree of cargo recruitment with respect to interaction domain concentration. We finally showcase compositional control of multiple proteins in a condensate to modulate protein interactions and tune metabolic flux.

## Results

### Protein-peptide Interaction Domains enrich protein cargo into synthetic condensates

To simplify the design of synthetic intracellular condensates, we utilized a one-component self-oligomerizing protein scaffold inspired by the optogenetic Corelet system ^40^. In this new design, clusters are assembled constitutively (i.e. not in a light-dependent manner), and are therefore termed “Forever Corelets” (FC). In the FC construct, the intrinsically disordered N-terminal region of the human FUS protein (IDR) is fused to human ferritin monomers, which self-assemble into oligomeric structures (‘Cores’) of 24 units decorated with 24 IDRs. The multimerization of these constructs amplifies the effect of the weakly interacting IDRs, resulting in protein condensate formation (Fig 1A, Top Panel). This approach is highly generalizable, allowing for the use of Cores other than ferritin and IDRs other than FUS^41,42^. Given the genetic versatility of *Saccharomyces cerevisiae*, including the ability to genomically integrate designer genes in defined copy numbers with well-characterized promoters of varying strength, we chose to develop our synthetic condensate framework in this yeast.

**Figure 1.**
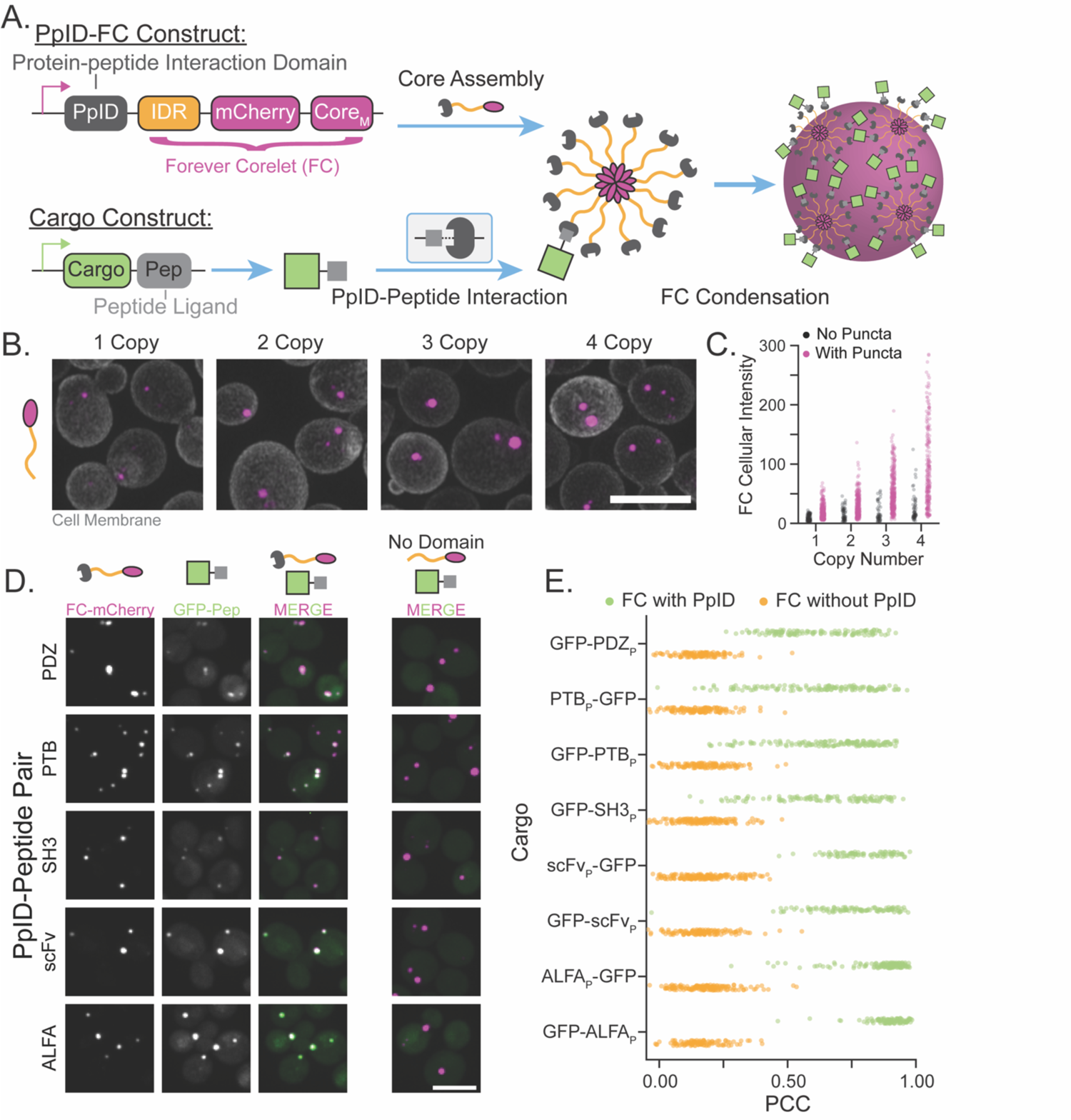
Protein-peptide Interaction Domains recruit cargo to synthetic condensates. A. Multimerization of IDRs through fusion to core monomers (Core_M_) yields constitutive puncta labeled with mCherry (Top Panel: PpID-FC Construct). Protein-peptide Interaction Domains are incorporated on the N-terminus of the FC construct to recruit cargo tagged with corresponding peptide (Bottom Panel: Cargo Construct). B. Representative FC puncta in cells expressing the FC construct at different copy numbers, as observed with Airyscan imaging. FC labeled with mCherry. Scale bar, 5 µm C. Mean FC fluorescence of cells without (black) and with (purple) puncta at each copy number of FC expression. (n>280 cells for each strain) D. Representative images of C-terminally tagged GFP-peptide cargo recruitment to PpID-FC assemblies (first three columns) as well as control strains without PpID fusion to FC (rightmost column). Merged images show PpID-FC in magenta, and GFP cargo in green. Scale bar, 5 µm E. Per-cell Pearson Correlation Coefficients (PCC) for strains expressing PpID-FC and the corresponding GFP-peptide fusion (Green) and control without PpID fusion to FC (Orange). Arranged in order of decreasing reported K_D_ (See Extended Data Table for values) (n>80 cells for each strain)

When the FC construct, labeled with the mCherry fluorescent protein, is expressed in *S. cerevisiae*, the cells constitutively show discernable puncta (Fig 1B), which exhibit larger sizes with increasing FC copy number, i.e., the number of FC genetic elements added to the cells (Extended Data Fig 1, Supplemental Note 1). Cells without discernible puncta exhibit very low mCherry signal, while increasing fluorescence intensity corresponds with an increasing fraction of cells exhibiting condensation (Fig 1C), typical of single component (i.e. excluding “solvent”) phase separating systems^9^; interestingly, the concentration dependent transition here is more broad than in the previously studied optogenetic counterparts^40,43^.

To harness FC condensates as a scaffold on which to programmably recruit non-phase separating proteins, we looked for small protein domains that, when incorporated into FC, could mediate specific scaffold-cargo interactions. We chose domains that bind small peptides of specific sequence, which by virtue of their size, minimize the risk of misfolding or inactivating cargo proteins. Domains of diverse origins were screened for recruitment of peptide-fused cargo to condensates (Supplemental Note 2). These included both synthetic and naturally occurring domains with reported dissociation constants (K_D_) on the order of 10^+2^ to 10^-5^ μM (Extended Data Table). We will refer to these domains as Protein-peptide Interaction Domains, or PpIDs. The larger peptide-binding domains are fused to the FC construct (Fig 1A, Top Panel), and the small peptides to which they bind are fused to either terminus of a cargo protein of interest (Fig 1A, Bottom Panel).

To screen these PpIDs, we used confocal microscopy to analyze the fluorescent signal overlap between GFP cargo and FC clusters. PpID-FC constructs were co-expressed with GFP cargo tagged with the corresponding peptide. Approximately half of the PpIDs (domain elements are indicated with the subscript ‘D’) we tested show prominent co-localization with their corresponding peptide-tagged (peptide elements are indicated with the subscript ‘P’) cargo (Fig 1D). These include: the PDZ domain from mouse a-Syn (PDZ_D_)^44^, the PTB domain from human neuronal X11 (PTB_D_)^45^, the SH3 domain from mouse Crk (SH3_D_)^46^, a scFv screened against yeast GCN4 (scFv_D_)^47^, and a Nanobody against the small ALFA peptide (ALFA_D_)^48^. While PTB_P_, ALFA_P_, and scFv_P_ allow for sufficient recruitment when fused to either terminus of the GFP cargo, PDZ_P_ and SH3_P_ only function as C-terminal fusions (Extended Data Fig 2, Supplemental Note 3). Quantifying the overlap using the Pearson Correlation Coefficient (PCC) (Fig 1E), we found that the PpIDs with the strongest binding to their peptide cargos (lowest K_D_ values, e.g. scFv and ALFA) exhibit more robust colocalization (higher PCC) compared to those pairs with weaker binding (higher K_D_ values, e.g. PDZ, PTB, and SH3).

### Degree of cargo enrichment increases with PpID-FC concentration

We next sought to evaluate the performance of this platform by a quantitative metric of cargo recruitment. One of the most intuitive and commonly used parameters of enrichment is the partition coefficient, defined as the ratio of concentration inside and outside the condensate. However, due to the diffraction-limited nature of typical puncta, it is difficult to accurately measure concentrations in the condensate (dense phase). Therefore, to quantify recruitment, we defined a related parameter we call ‘recruitment’, or R, that reflects the fraction of cargo enriched in the condensate through direct interaction with PpIDs:

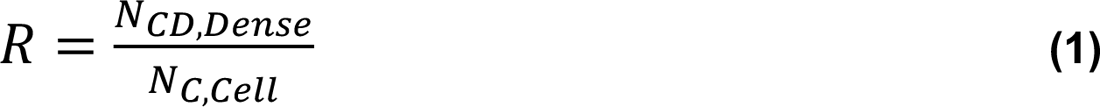

where N_CD,Dense_ is the number of cargo molecules (C) bound to a PpID (D) in the dense phase and N_C,Cell_ is the total number of cargo molecules in the whole cell system (Fig 2A). Recruitment can be determined experimentally based on the fluorescence intensities of GFP used as cargo and mCherry-tagged PpID-FC in the whole cell and dilute phase regions (Supplemental Note 4.2). R ranges from nearly 0, in systems that have no cargo recruitment, as in negative controls, and increases towards 1, which would reflect complete cargo recruitment. When applied to images of yeast strains expressing PpID-FC and GFP tagged with the PpID cognate peptide, we observe a significant increase in R for each PpID compared to controls of FC expression without PpID (Extended Data Fig 3).

**Figure 2.**
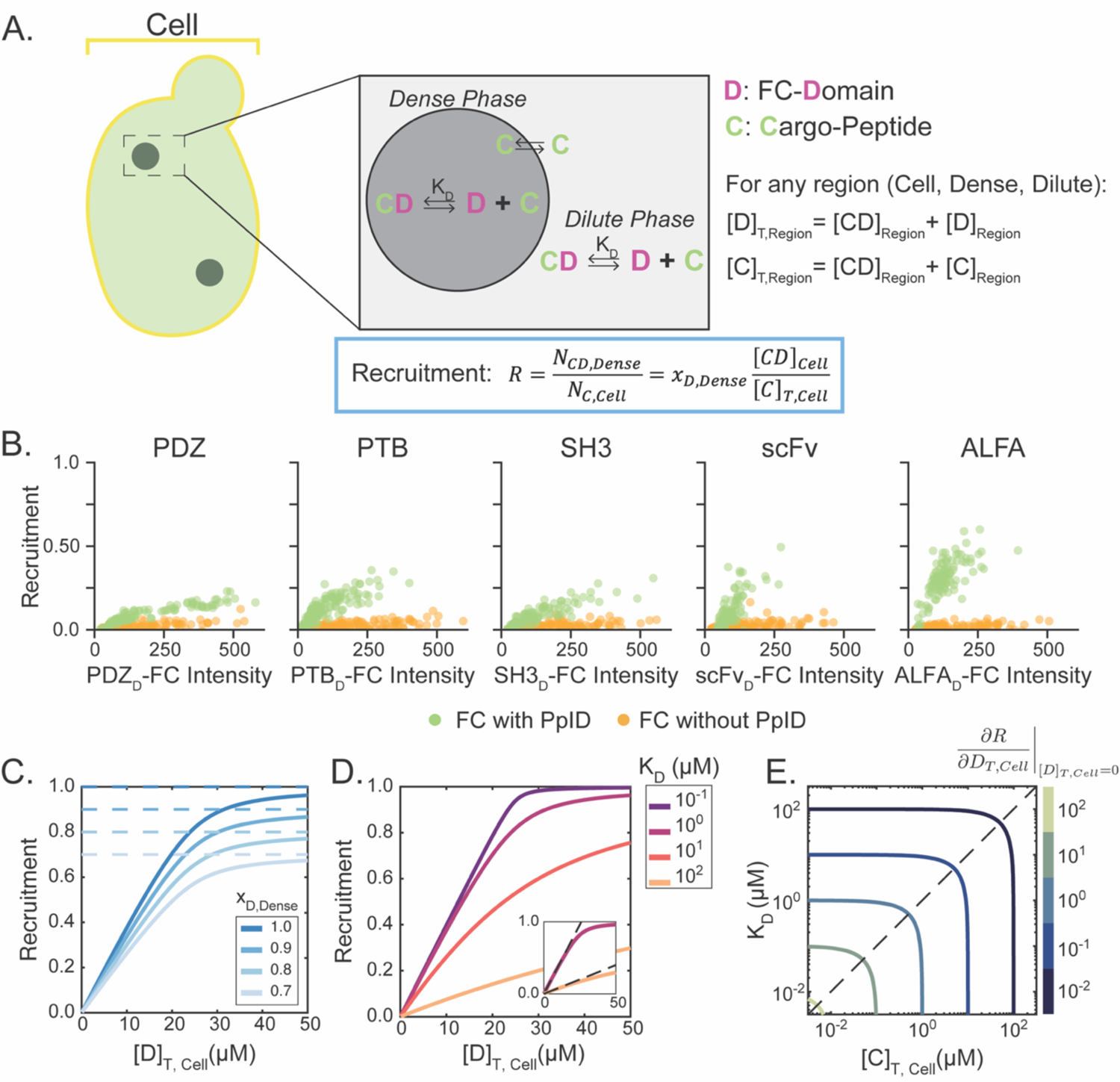
Recruitment increases with PpID-FC concentration. A. Recruitment is defined as the fraction of cargo molecules in the cell that are recruited into condensates via PpID-cargo interactions (when cargo is peptide-tagged). C: cargo, D: PpID B. C-terminally tagged GFP cargo recruitment per cell with respect to PpID-FC mean cellular fluorescence intensity. Plots arranged in order of decreasing reported K_D_ (See Extended Data Table for values) (n>100 cells for each strain) C. Analytical solutions for recruitment as a function of [D]_T,Cell_ with constant [C]_T,Cell_ = 25 μM, K_D_ = 1 μM, at various values of x_D,Dense_. Dashed lines indicate the maximum for each condition, as given by x_D,Dense_ D. Analytical solutions for recruitment as a function of [D]_T,Cell_ with constant [C]_T,Cell_ = 25 μM, x_D,Dense_ = 1. (Inset) Dashed line shows initial slope for the cases of K_D_ = 1μM and 100μM, as given by equation (3). E. Contour plot of equation (3) for the initial slope of R with respect to [D]_T,Cell_ for x_D,Dense_ = 1. Dashed line shows [C]_T,Cell_ = K_D_ curve

Interestingly, while PDZ (which has the highest K_D_) shows the lowest cargo recruitment and ALFA (with the lowest K_D_) shows the highest R, this trend with K_D_ does not hold well for all cases. In particular, scFv shows bulk recruitment similar to the PDZ, PTB, and SH3, despite having a K_D_ similar to ALFA (Extended Data Fig 3). We considered that this lack of correspondence between K_D_ and R might be due to differences in expression levels when different PpIDs were utilized. Indeed, over the range of PpID-FC expression levels assayed in these cells, we found that R monotonically increases with the PpID-FC (mCherry) fluorescence intensity, becoming increasingly steep with decreasing K_D_ (Fig 2B). This is consistent with intuition: as more PpID-FC is expressed, more domains are present to bind peptide-tagged cargo and enrich them in the FC condensate. Furthermore, as K_D_ decreases, a larger fraction of domains exist in a bound state, resulting in increased R at a particular PpID-FC concentration.

While we observe a striking recruitment-dependence on PpID-FC fluorescence level (Fig 2B), R shows essentially no dependence on cargo concentration (Extended Data Fig 4). This lack of dependence with cargo concentration is, at first glance, unexpected. From a protein-binding standpoint, increasing cargo concentration should increase the concentration of the bound cargo-PpID population. However, increasing cargo concentrations could also lead to excess unbound, and therefore unrecruited, cargo in the dilute phase of the cell. These considerations underscore the non-intuitive effects at play, pointing to the need for a quantitative modeling approach.

### Recruitment can be modeled by equilibrium binding

To better understand the system parameters affecting recruitment, we developed a simple analytical framework to describe cargo recruitment. Building from earlier models of client recruitment to condensates^35,49,50^, we developed a mass-action model of cargo binding equilibrated over the dilute and dense phases of PpID-FC. In this model, we consider the PpID-FC as partitioning domains, D, between the dense and dilute regions of the total cell system (Fig 2A). Although the binding and recruitment of cargo is specifically mediated by PpID-peptide interaction, for simplicity we will hereon refer to domain-cargo interactions as the basis for recruitment. Cargo molecules, partition between the phases via interactions with domains as dictated by the equilibrium dissociation constant, K_D_, which is assumed to be equal in the dense and dilute phases. Furthermore, we assume that the unbound cargo can freely diffuse between regions, so that the concentration of this population is equal in the entire cell. With these assumptions, the bound population, [CD]_Cell_, can be solved analytically in a manner analogous to receptor-ligand binding (Supplemental Note 4.3). Critically, R can be rewritten in terms of the cellular concentration of the bound cargo-domain assembly, [CD]_Cell_ (Supplemental Note 4.3):

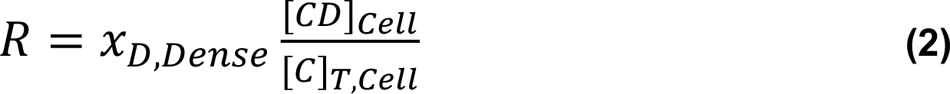

Here *x_D,Dense_* = *N_D,Dense_*/*N_D,Cell_* is the ratio of domain molecules in the dense phase to the whole cell, which depends on the condensate volume fraction, and density (Supplemental Note 4.2, 4.3). [C]_T,Cell_ is the cellular concentration of all cargo populations (subscript T: total, bound and unbound)(Fig 2A).

Expressing recruitment as shown in equation (2) illustrates that the condensate system imposes an upper limit on R, as expressed by x_D,Dense_ (Fig 2C). Even in a system with complete binding of cargo to domains, recruitment is limited by the fraction of domains that exist in the dilute phase (outside of the condensates). Indeed, measured recruitment in systems with increasing domain concentration in the dilute phase decreases R in a manner directly proportional to x_D,Dense_ (Extended Data Fig 5). Therefore, the observed recruitment is the product of the extent of domain enrichment in the dense phase, as determined by the condensate properties (i.e. through x_D,Dense_), and the fraction of those domains that are bound to cargo, which is dictated by the interplay of K_D_, [D]_T,Cell_ and [C]_T,Cell_.

We observe that K_D_ scales the recruitment response to domain concentration, but only when K_D_ is equal or greater in magnitude than cargo concentration (Fig 2D). For the highest K_D_ considered, R increases over a large range of domain concentrations. As K_D_ decreases, the model shows a steeper increase in recruitment with respect to domain concentration, replicating the shift in R response observed experimentally between the PpIDs of various binding strengths (Fig 2B). The effect of K_D_-scaling appears to saturate, as lower values of K_D_ are considered. This behavior can be understood as an interchanging effect between K_D_ and [C]_T,Cell_, as captured in the initial slope of the recruitment response (Supplemental Note 4.3; Fig 2D, insert):

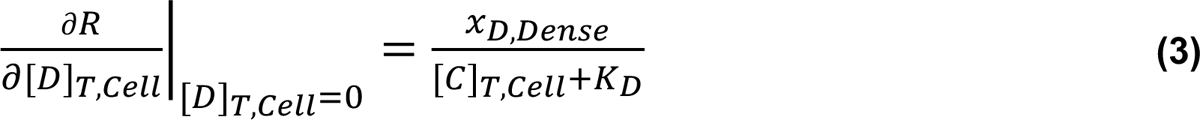

From this expression, it can be seen that the values of [C]_T,Cell_ and K_D_ symmetrically affect the initial trajectory of the recruitment dependence on [D]_T,Cell_, such that as K_D_ decreases below [C]_T,Cell_, the latter begins to dictate the response (Fig 2E), and vice versa.

### KD and cargo concentration dictate the recruitment response to PpID concentration

Equation (3) suggests an interplay between K_D_ and [C]_T,Cell_, which dictates how R responds to [D]_T,Cell_. We therefore considered the limiting cases where either K_D_ or [C]_T,Cell_ is significantly greater than the other to develop an understanding of the nature of recruitment (Supplemental Note 4.4). For [C]_T,Cell_ << K_D_, R takes the form of a Langmuir equation (Fig 3A), which shows no dependence on cargo concentration:

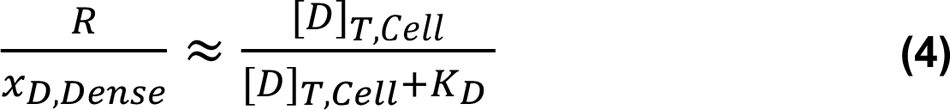

**Figure 3.**
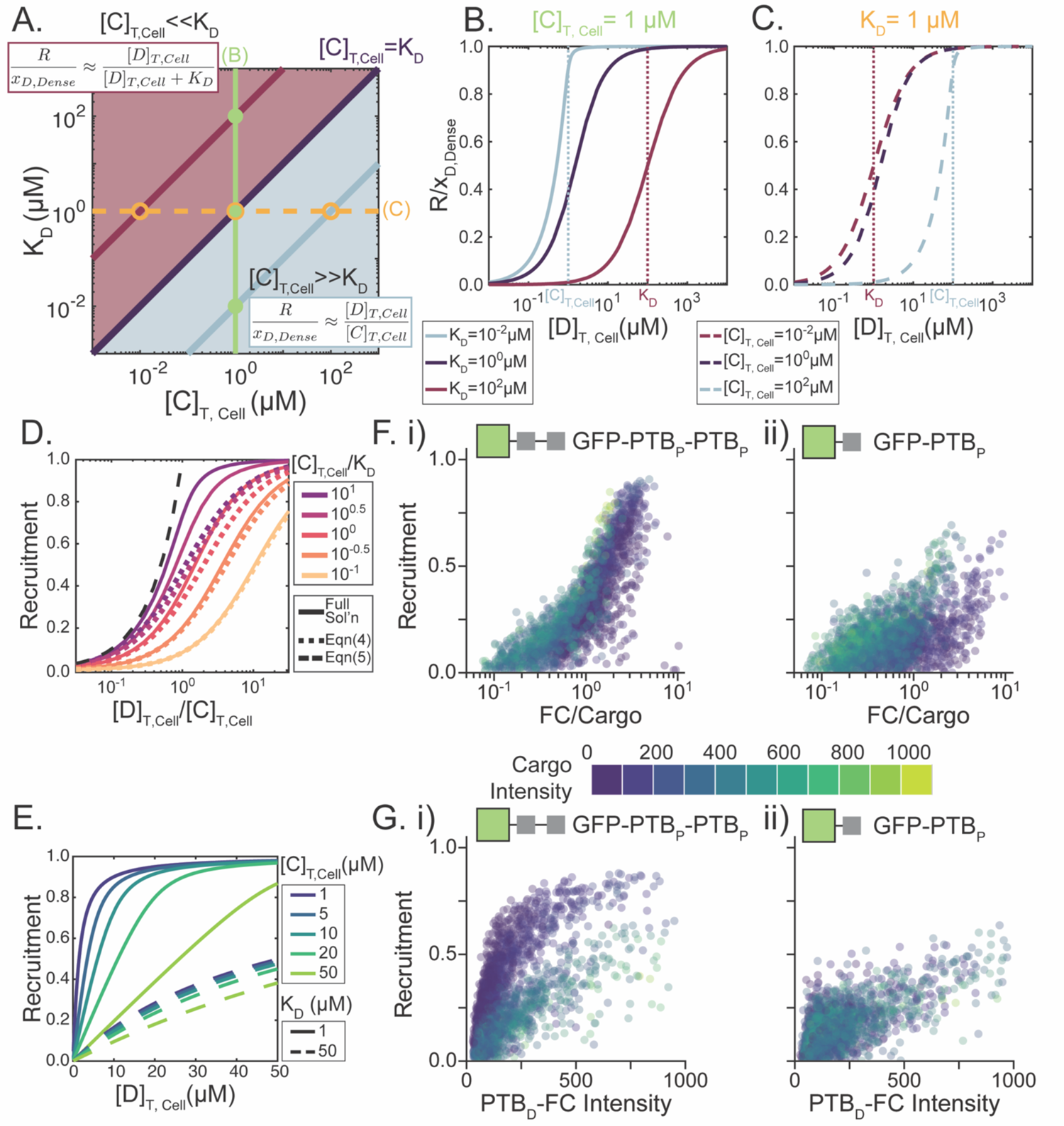
K_D_ and cargo concentration dictate recruitment response to PpID concentration. A. The K_D_/[C]_T,Cell_ ratio dictates the form of R response to [C]_T,Cell_. Green and orange lines respectively indicate K_D_ and [C]_T,Cell_ values while maroon and powder blue curves respectively indicate [C]_T,Cel_<<K_D_ and [C]_T,Cell_>>K_D_ conditions considered in (B&C). B. Recruitment response to [D]_T,Cell_ with constant [C]_T,Cell_=1 μM in the [C]_T,Cell_>>K_D_, [C]_T,Cell_=K_D_, and [C]_T,Cell_<<K_D_ regimes. Dotted lines indicate the value of [C]_T,Cell_ in the case of [C]_T,Cell_>>K_D_ and K_D_ in the case of [C]_T,Cell_<<K_D_. x_D,Dense_ = 1 for all curves C. Recruitment response to [D]_T,Cell_ with constant K_D_=1 μM in the [C]_T,Cell_<<K_D_, [C]_T,Cell_=K_D_, and [C]_T,Cell_>>K_D_ regimes. Dotted lines indicate the value K_D_ in the case of [C]_T,Cell_<<K_D_ and [C]_T,Cell_ in the case of [C]_T,Cell_>>K_D_. x_D,Dense_ = 1 for all curves D. Analytical solutions for recruitment (solid lines) as a function of [D]_T,Cell_/[C]_T,Cell_ for various [C]_T,Cel_/K_D_ regimes for x_D,Dense_ = 1. K_D_-dominated regime approximations (equation (4)) shown as dotted lines for each condition as color coded, while the complete binding approximation (equation (5)) is shown as a black dashed line. E. Analytical solutions for recruitment as a function of [D]_T,Cell_ at various K_D_ and [C]_T,Cell_ values with x_D,Dense_ = 1. F. Per-cell recruitment with respect to [D]_T,Cell_/[C]_T,Cell_ (based on normalized mean cellular intensity values of PTB_D_-FC and GFP Cargo) with color-coded bins for cargo intensity for (i) GFP-PTB_P_-PTB_P_ and (ii) GFP-PTB_P_ cargo G. Per-cell recruitment with respect to PTB_D_-FC mean cellular fluorescence intensity with color-coded bins for mean cellular cargo intensity for (i) GFP-PTB_P_-PTB_P_ and (ii) GFP-PTB_P_ carg For F&G) Color bar for cargo cellular intensity considers normalization of the relative levels of fluorescence intensity of mCherry and GFP (n>2000 cells for both (i) and (ii))

In this regime, R reaches half maximum when [D]_T,Cell_ is equal to K_D_, as is typical for the Langmuir equation^51^ (maroon curves, Fig 3B,C). We will refer to this regime as the ‘K_D_-dominated regime’.

Conversely, for [C]_T,Cell_ >> K_D_, R no longer shows the effect of the K_D_-defined binding equilibrium (Fig 3A):

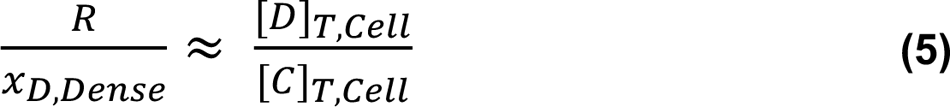

which captures the recruitment behavior until R/x_D,Dense_ = 1. In this limit, it is expected that all domains are bound to cargo molecules and therefore R increases in a manner dependent on the ratio of total domain and cargo concentrations. Therefore, R approaches its maximum value when [C]_T,Cell_ is equal to [D]_T,Cell_ (teal curves, Fig 3B,C). In this regime, [D]_T,Cell_ = [CD]_Cell_, so we therefore refer to this case as the ‘complete-binding regime’.

These theoretical considerations imply that the ratio [C]_T,Cell_ / K_D_ dictates which behavior will predominate, and can provide insight into the expected recruitment behavior, even away from the limiting cases considered above. In particular, when [C]_T,Cell_ / K_D_ is above unity, R is well approximated by equation (5), while cases of [C]_T,Cell_ / K_D_ below unity show behavior predicted by equation (4) (Fig 3D). With this view, we rationalize that the apparent lack of cargo concentration sensitivity observed experimentally (Extended Data Fig 4) suggests that our cargo expression levels (relative to K_D_) position cells in the K_D_-dominated regime. Indeed, the analytical model showcases a much wider range in R response for varying cargo concentrations in the case of [C]_T,Cell_ > K_D_ (Figure 3E, solid lines), compared to [C]_T,Cell_ < K_D_ (Figure 3E, dashed lines). Nevertheless, the experimental measurements of R were performed at a single cargo expression level, which translates to a lesser range in measured concentrations than that obtained for the FC component (Extended Data Fig 4).

To verify the interpretations and applicability of the model framework outlined above, we sought to explore a wider range of [C]_T,Cell_ and K_D_ experimentally. Instead of using various PpIDs to sample various binding strengths, we chose to focus only on PTB and use different cargo architectures to lower the effective K_D_. PTB shows an intermediate binding affinity amongst the PpIDs tested (Fig 2B), away from the complete-binding regime behavior observed with ALFA and scFv. To lower the apparent K_D_ of the PpID-cargo interaction, we increased the peptide valency on our GFP cargo by incorporating a tandem repeat of PTB_P_. Multivalency in ligand-receptor systems is broadly seen to lead to ‘avidity’ or a decrease in the effective K_D_^35,52,53^, allowing for higher fractional binding (Extended Data Fig 6). We therefore engineered yeast strains to express PTB_D_-FC and cargo with one or two PTB-binding peptides (GFP-PTB_P_ or GFP-PTB_P_-PTB_P_) at various levels using promoters of different strengths and multiple gene copy numbers (Extended Data Fig 7).

The two cargo architectures show a change in [C]_T,Cell_ / K_D_ regime that supports the predicted relative sensitivity to cellular cargo concentrations. For the case of GFP-PTB_P_-PTB_P_, when recruitment is plotted against [D]_T,Cell_ / [C]_T,Cell_, the response curves collapse for the range of cargo intensities observed (Fig 3F(i)), indicative of [C]_T,Cell_-scaling in the ‘complete-binding regime’. Likewise, R plotted against PTB_D_-FC intensity showcases distinct sensitivity to the concentration of this cargo (Fig 3G(i)). Conversely, for GFP-PTB_P_ cargo, the cargo intensity normalization does not collapse the recruitment response (Fig 3F(ii)), and the data instead match the model curves for increasing cargo concentrations in the K_D_-dominated regime (Fig 3D). Consistently, a lack of R sensitivity to cargo concentration is still observed for the monomeric peptide tag (Fig 3G(ii)), even with the increased range of cellular concentrations achieved in these experiments (Extended Data Fig 7). The consistency of the experimental outcomes with the well-defined theory of equilibrium-binding underscores that PpID-based cargo recruitment to cargo represents an experimental framework for rational design of functionalized condensates.

### PpIDs enable stoichiometric control of multiple cargo in a condensate

With a clear understanding of the characteristics of cargo recruitment to condensates, we verified that these properties extend to simultaneous recruitment of multiple cargo proteins. We first confirmed that the PpIDs are orthogonal and do not exhibit cross interactions with peptides other than their cognate partner. We imaged yeast cells expressing each of the five previously identified PpIDs on mCherry-tagged FC combinatorially with GFP cargo fused to each cognate peptide on either terminus (where applicable). For each pair, we quantified the colocalization of the mCherry and GFP signals with a cell-averaged Pearson Correlation Coefficient (Fig 4A). We only see enhanced correlation values along the diagonal, in strains where the FC construct being expressed contains the PpID known to interact with the peptide fused to the co-expressed GFP cargo. This confirms that all the tested PpIDs are orthogonal, which enables independent recruitment of multiple cargo proteins with stoichiometric control.

**Figure 4.**
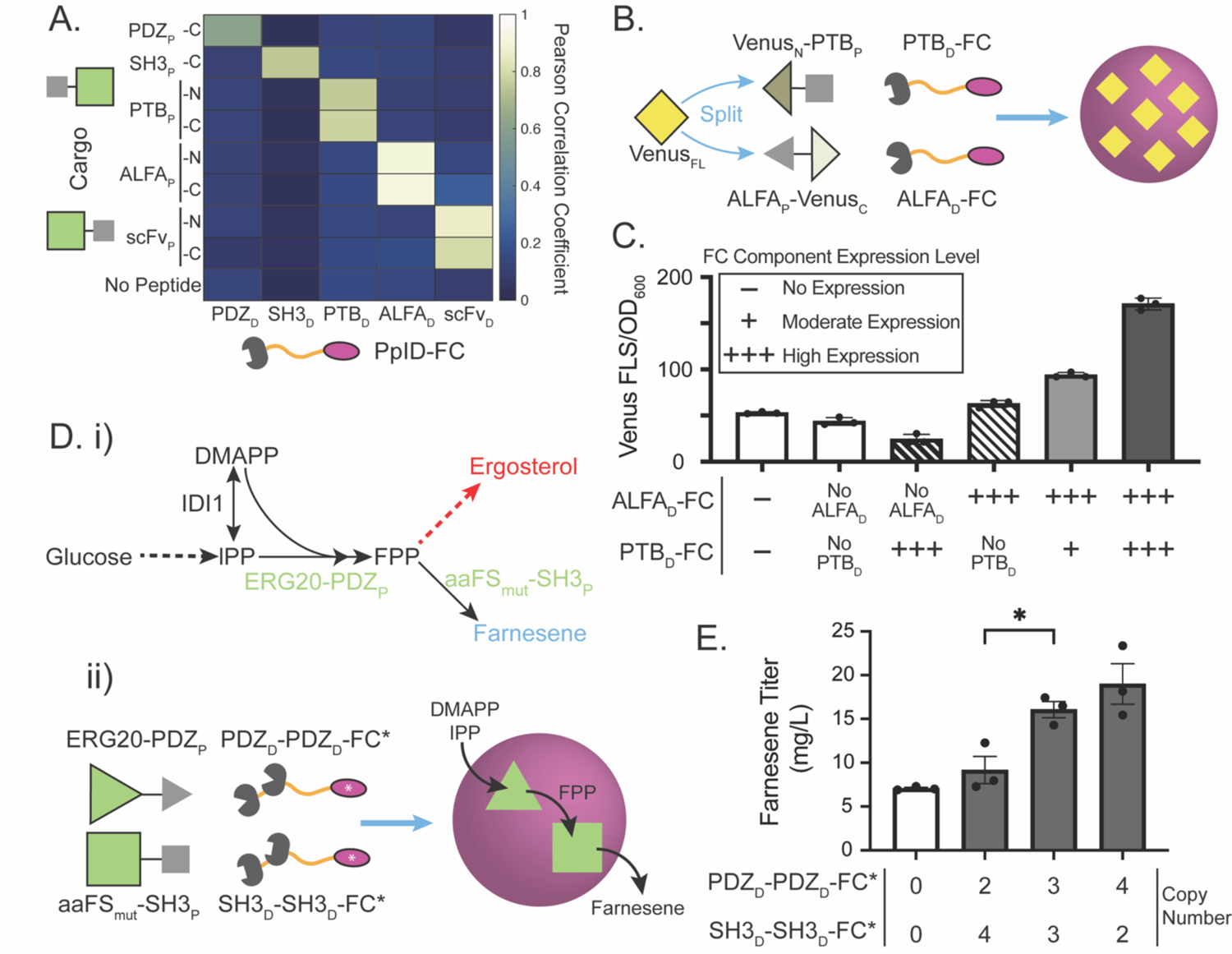
Multiple PpIDs can be used simultaneously to tune cargo stoichiometry. A. Matrix of Pearson Correlation Coefficients between Cargo and FC signal for each PpID-peptide pairing B. For vPCA, PTB_D_ and ALFA_D_ were used to recruit two fragments of Venus to FC clusters to enable refolding of Venus and functional fluorescent output C. Bulk OD-normalized Venus fluorescence in cultures of yeast expressing various PpID-FC combinations in a vPCA parent strain, as measured by a plate reader. Bar pattern indicates PpID expression: white, no PpIDs expressed; striped, only one PpID expressed; gray both PpIDs expressed, darkening with increased expression. Mean and standard error shown for n=3 technical replicates for each strain. D. (i) Engineering farnesene production in yeast is accomplished by the expression of aaFS_mut_ to divert FPP from the native ergosterol biosynthesis pathway. The enzymes targeted for condensate recruitment are labelled in green. (ii) Double PpID fusions of SH3_D_ and PDZ_D_ to I3-01-based FC (FC*, Supplemental Note 6) are used to recruit aaFS_mut_ and ERG20, respectively, to condensates with the goal to streamlining farnesene production E. Farnesene production as a function of different stoichiometries of SH3_D_ and PDZ_D_ in the FC* condensates. The copy number of SH3_D_-SH3_D_-FC* and PDZ_D_-PDZ_D_-FC* are indicated below each column. Control with no domains expressed shown with white bar. Mean and standard error shown for n=3 biological replicates for each strain. *: p-value<0.05, Statistics from two-sided t-test

To examine the tunable recruitment of multiple cargo species in a condensate, we first employed a Venus Protein-fragment Complementation Assay (vPCA). In vPCA, the Venus fluorescent protein is expressed in two different fragments (Venus_N_ and Venus_C_, Fig 4B), which bind and refold to make an active Venus protein when brought into close proximity, such as when fused to interacting proteins^54,55^. Leveraging the observation that the degree of protein recruitment markedly increases with increasing expression levels of the corresponding PpID-FC, we can use different PpID pairs to independently tune the recruitment levels of each fragment. Specifically, the Venus N- and C-fragments were fused with PTB_P_ and ALFA_P_, respectively, and expressed using the moderate strength HHF2 promoter in all strains (Fig 4B). We expressed ALFA_D_-FC at equal levels using the strong CCW12 promoter, and varied expression of PTB_D_-FC using constitutive promoters of different strengths. When using the moderate HHF2 promoter to express PTB_D_-FC, there is a 2-fold increase in Venus fluorescence signal over controls without recruitment. Furthermore, the output is over 3-fold higher than the controls when PTB_D_-FC expression was further increased using the stronger TDH3 promoter (Fig 4C). Strains expressing FCs without PpIDs or only one of the PpIDs, largely decreased fluorescence compared to a strain without any FC expression (Supplemental Note 5). This confirms that FCs containing both PpIDs are required to recruit both Venus fragments to the FC condensates, and that the proportion of each recruited fragment can be manipulated by varying the level of expression of each FC-PpID.

We next tested whether PpID-FC system can provide precise control of more complex biological processes recruited to the FC condensate, such as flux regulation through a branchpoint in metabolic pathway. Clustering metabolic enzymes that produce and consume intermediates can mitigate flux to byproduct formation catalyzed by unclustered competing pathways^30,56^. This is primarily achieved through a mass action effect, which competes with the diffusion of the intermediate out of the enzyme cluster^56^. Therefore, the relative rates of reaction of each recruited enzyme, which are dictated by the enzyme concentrations in the cluster, play a crucial role in optimizing the metabolic flux through a clustered pathway. We thus hypothesized that the tunability afforded by our orthogonal PpID system is ideal to control engineered metabolic pathways clustered in synthetic organelles.

To test this hypothesis, we used our PpID-based synthetic organelles to compartmentalize the biosynthesis of β-farnesene, a sesquiterpene that can be used as renewable jet fuel^57^; notably, the pathway competes for farnesene pyrophosphate (FPP) with the native biosynthesis of ergosterol, an essential lipid^58^ (Fig 4D). We clustered a farnesene synthase that was previously engineered for enhanced activity (aaFS_mut_)^59^ in synthetic FC-like condensates (FC*, Supplemental Note 6) with ERG20, the immediately preceding enzyme that catalyzes the coupling of dimethylallyl pyrophosphate (DMAPP) with two molecules of isopentenyl pyrophosphate (IPP) to produce FPP, the substrate of farnesene synthase^58^ (Fig 4D). While keeping enzyme expression and the total copy number of FC* the same, we titrated the stoichiometry of ERG20-PDZ_P_ and aaFS_mut_-SH3_P_ recruited to condensates by varying the copy number of the FC* constructs fused to a tandem repeat of the corresponding PpID (PDZ_D_-PDZ_D_-FC* and SH3_D_-SH3_D_-FC*, respectively) We observed that higher levels of PDZ_D_-PDZ_D_-FC* (and therefore ERG20) in the cluster, relative to SH3_D_-SH3_D_-FC* (and therefore aaFS_mut_), significantly increase farnesene production (Fig 4E). This makes sense, as increasing the availability of FPP in the clusters channels more of this precursor to β-farnesene production. The optimal stoichiometry of recruited enzymes (their ratios in the condensate) will depend on their relative enzymatic rates, which can be time-consuming to accurately measure intracellularly. Therefore, having a method to systematically screen different ratios of recruited enzymes greatly simplifies the optimization of metabolic pathways compartmentalized in synthetic organelles.

## Discussion

In this manuscript, we set out to develop a strategy for programming protein recruitment to intracellular condensates to decouple condensate formation and functionalization, to enable tunability, and to achieve orthogonality in multi-cargo recruitment. The use of PpIDs is at the heart of satisfying these goals (Fig 5). In using binding domains, we separate the relative expression of condensate and cargo constructs. The freedom to separately control the expression level of these components imparts tunability to the degree of cargo recruitment (Fig 5A). The flexibility of this design is further expanded by using different orthogonal PpIDs with varying peptide-binding affinities. As each PpID shows no crosstalk, these strategies can be coupled to provide flexible programmability to the recruitment of multiple cargo proteins to synthetic condensates (Fig 5B,C).

**Figure 5.**
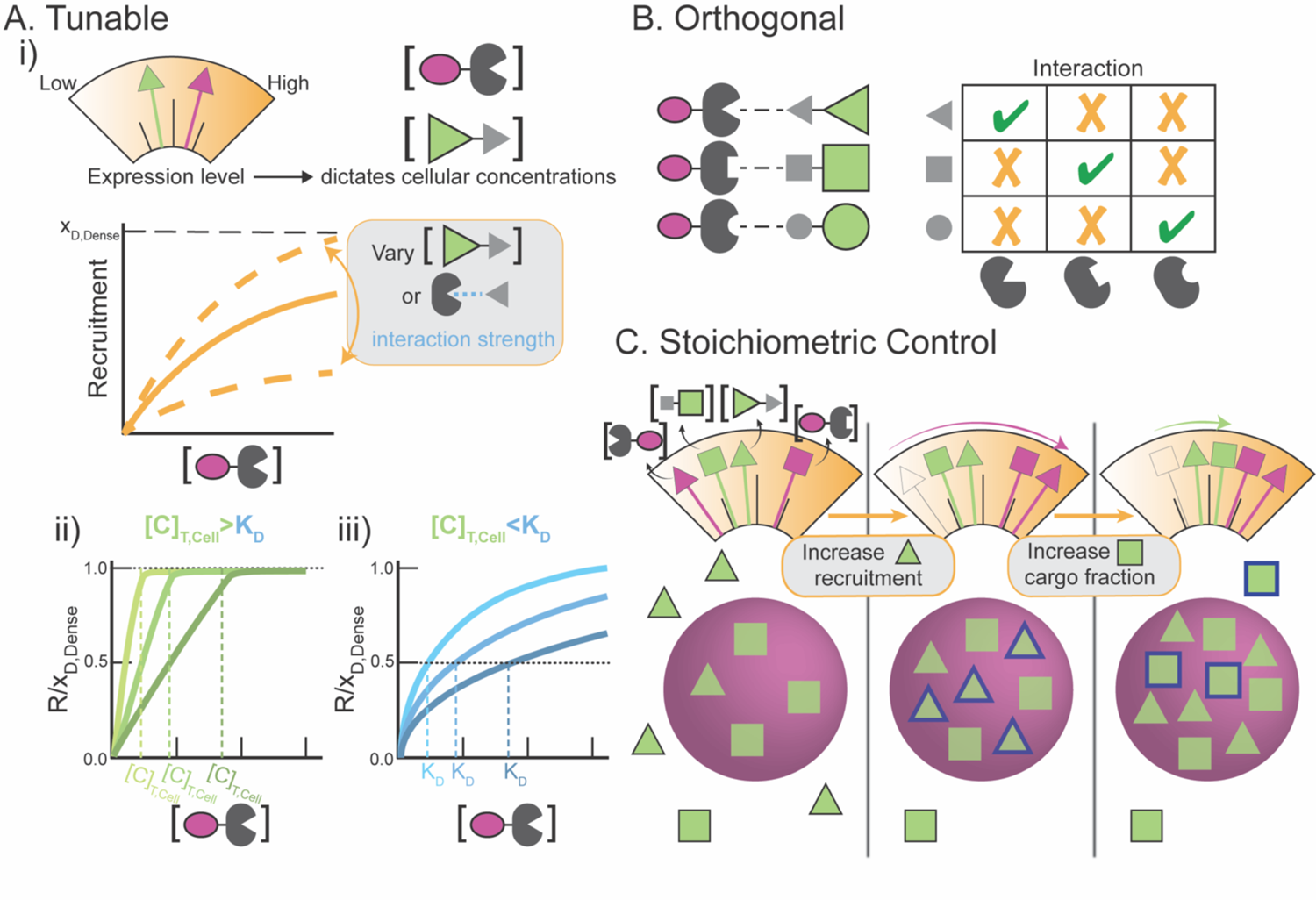
PpID-based cargo recruitment enables predictable engineering of synthetic condensates. _A._ The PpID system is tunable: (i) Expression levels of condensate-PpID (purple oval, any condensate forming component, fused with PpID) and cargo (green triangle, tagged with peptide) components can be independently controlled to tune cargo recruitment response. Recruitment increases as a function of PpID concentration to a maximum equal to x_D,Dense_ (orange curves). Changes in cargo concentration and domain-peptide interaction strength tunes the response of recruitment to domain concentration (dashed orange lines). The behavior of these responses can be generalized into two cases: (ii) ‘Complete Binding’ when cargo concentration is greater than the K_D_ of the PpID-peptide pair, and (iii) ‘K_D_-dominated’ when cargo concentration is lower than the K_D_ of the PpID-peptide pair; for (ii) and (iii) various exemplar curves for each regime is illustrated schematically where darker color corresponds to states of (ii) increasing cargo concentration and (iii) increasing K_D_ B. The PpID system is orthogonal: A library of orthogonal interaction domains enable flexible recruitment of multiple cargo species C. In tandem, these design features of the PpID system enable multi-cargo stoichiometric control inside synthetic condensates. Degree of cargo recruitment is largely dictated by expression level of the corresponding condensate-PpID construct, while relative levels of cargo protein are tuned by their expression level. Changes in cargo composition are highlighted in blue.

Based on the ability to cast recruitment in an equilibrium-binding framework, one can experimentally tune the nature of cargo recruitment with respect to PpID-FC expression by selecting particular cargo concentrations and PpIDs with specific K_D_ values (Fig 5A(i)). As evident from the initial slope of R with respect to [D]_T,Cell_ (equation (3)), when both K_D_ and cargo concentration are low, a rapid increase in recruitment with respect to PpID concentration is observed. Moreover, low K_D_ compared to cargo concentration leads to a linear increase in recruitment (Fig 5A(ii)), which can be approximated to complete domain binding. Conversely, as K_D_ increases relative to cargo concentration, recruitment shows a more gradual increase scaled by K_D_ (Fig 5A(iii)), which is largely insensitive to cargo concentration, making it ideal to enable more predictable recruitment in cases of unsteady or unknown cargo concentration.

Modular design of synthetic organelles for engineering applications is a widely-appreciated goal^38,39^. Our platform meets this challenge by being highly modular and programmable to enable future “plug-and-play” capabilities which can be readily incorporated for a variety of distinct synthetic organelles for various applications. While our study focuses on FC condensates, the PpIDs used and the principles derived here are extendable to other clustering approaches. This could provide a powerful strategy to tune recruitment by adjusting x_D,Dense_ via selection of different condensate systems (Extended Data Fig 5). The x_D,Dense_ could also be tuned through subcellular spatial activation of optogenetic condensate systems, providing temporal control over condensate assembly and cargo recruitment^40,42^. Additionally, chemical or optogenetic transcriptional control of cargo and PpID-FC components could also enable in situ programming of recruitment^60^.

Another future direction will focus on expanding the library of PpIDs applicable to functionalize our designer condensates. As longer peptides present more moieties with which to interact, there is a general correlation between peptide size and binding affinity among PpIDs. Indeed, recruitment has been shown to increase through the use of much lower K_D_ domains, which also tend to bind longer length peptides^26,27^. Nonetheless, we and others have observed a detrimental effect on protein activity as fusion size increases^61^. As a result, we here focused on domains that bind small ligands, many of which also have relatively large K_D_. Given this tradeoff, characterizing a larger library of domains that offer a range of attributes, whether it be a particular binding affinity or ligand length, would prove useful. While nanobodies and scFv’s can defy the K_D_-ligand length trend, we and others have observed intracellular instability of some of these domains^62^ (Supplemental Note 3). Excitingly, de novo design of novel PpIDs could enable a range of peptide lengths and binding affinities useful for this application^63^.

PpID-based functionalization of synthetic condensates thus provides the basis for the continued adoption of membraneless organelles as hubs for synthetic biology^6,23,24^. In particular, recruiting metabolic pathways to condensates can divert flux of intermediates away from competing pathways^30–34^. Indeed, the relative stoichiometry and degree of recruitment of enzymes can significantly impact the resulting product formation^56^. The PpID framework outlined above provides a powerful degree of control over these parameters, enabling metabolic engineering applications that finely tune the system composition to optimize performance. Such tunability will prove particularly empowering when specific relative rates of flux to different products is desirable^64^. The platform we present here to systematically functionalize and control the composition of synthetic organelles will thus facilitate the use of biomolecular condensates in a wide range of synthetic biology applications.

## Methods

### Molecular Biology

#### Assembly of DNA constructs

All plasmids were cloned into the standardized pJLA vector series^65^. FC were constructed via the M22 and M23 Corelet plasmids^40^ and assembled via Gibson assembly. The NotI and NheI restriction sites upstream of the ORF in pJLA vectors were used to add PpIDs to the N-terminus of FC. In the pJLA vector series, the coding sequence is flanked upstream by NotI-KOZAK-NheI and downstream by XhoI-STOP codon. These restriction sites were leveraged to add the PpID ligand tags to the GFP cargo. All such modifications were made via oligo annealing to create the ligand with a (G4S)2 flexible linker and corresponding sticky ends for ligation. Other proteins were tagged with ligands by replacing GFP via NheI/NhoI cutsites. In most cases, (PpID)-FC constructs were subcloned via restriction/ligation into His3 locus integration plasmid, while cargo constructs were similarly subcloned into Leu2 integration plasmid, both for chromosomal integration into the yeast genome. Multigene cassettes were constructed using the sequential gene insertion sites built into the pJLA vectors. All oligonucleotides were synthesized by Genewiz and Integrated DNA Technologies. All enzymes used for construct generation are from New England Biolabs and Thermo Scientific. All oligos were ordered from Integrated DNA Technologies.

Genes for the PpIDs were synthesized by Synbio Technologies, flanked by NotI and NheI for cloning on to the FC Construct. Sequences for the PpIDs and cognate peptides are from the references listed in Extended Data Table 1. Promoters and terminators not originally in the pJLA series were taken from The MoClo-YTK plasmid kit, a gift from John Dueber (Addgene kit # 1000000061). The constructs encoding sequential incorporation of serine in place of tyrosine in the FUS IDR was based on previously published sequences^66^. The following coiled-coil domains were used to form dimeric and trimeric GFP cargo: BCR antiparallel homodimer^67^ and 5L6HC3_1 Untwisted homotrimer^68^. The sequence for aaFS_mut_ was based sequence ‘FS_C_8’ on a reported screen of mutations in Artemisia annua β-farnesene synthase^59^. All plasmids used in this study can be found in Supplemental Table 1.

### Yeast Strain Development and Protocols

All *Saccharomyces cerevisiae* in this study are of CEN.PK2-1C origin. All transformations were carried out using standard lithium acetate protocol^69^ using DNA linearized with PmeI or SwaI (New England Biolabs) to produce genomically integrated cells; in a few cases plasmids (CEN or 2μ) were directly used to introduce genetic material to the cells. Cells were grown in synthetic complete media with 2 % glucose (SC), at 30°C and shaking at 200 RPM. Growth for experimentation was carried out in 1 mL liquid cultures in 24 well plates (CELLTREAT). Knockout media (corresponding to auxotrophic markers restored in the corresponding strain) was always used for initial outgrowth of colonies from agar plates. When CEN or 2μ plasmids were used, cells were always maintained in Uracil knockout media to retain the plasmid. All yeast strains used in this study can be found in Supplemental Table 2.

### Microscopy

#### Sample Preparation

Except where stated otherwise, all imaging was performed on yeast harvested during early exponential phase. Briefly, a streak of multiple colonies of a yeast strain were grown overnight in selective media (typically SC-His-Leu) to saturation. The next day, the saturated overnight culture was diluted 20-40 fold in 1 mL fresh SC media. Following growth for 4-6 hours, cells are verified to be in early exponential growth, as defined by having OD_600_ in the range of 1.5 – 3 as approximated with a plate reader (TECAN). Cells were diluted to an OD_600_ of 1 in PBS (gibco); 100 μL of this dilution was added to Concanavalin A (Sigma-Aldrich) treated 96-well glass plate (Cellvis), with one strain per well. Cells were treated with Membrite Fix 640/660 membrane dye (Biotium) following manufacturer’s protocol. Typically, cells were then fixed with 4% paraformaldehyde (Electron Microscopy Science), diluted in PBS, for 15 minutes. Cells were washed 3 times with PBS and were imaged in fresh PBS. For these samples, we have observed fixation to faithfully represent what is observed in cells imaged live.

For assaying cells at different stages of growth, cells were initially prepared as stated above. Cells were harvested and prepared for imaging as previously described at either 8, 24, or 48 hours of growth (instead of 5 hours). For the case of 48 hours, cells were pelleted and resuspended in fresh media at the 24 hour time point.

#### Spinning Disk Confocal Microscopy

For all microscopy-based experiments, unless specified otherwise, confocal images were collected using a Yokogawa CSU-X1 spinning disk with a 100x, NA 1.49, Apo TIRF oil immersion objective on a Nikon Eclipse Ti body and Andor DU-897 EMCCD camera, all controlled via Nikon Elements 5.02. Protein fusions containing GFP and mCherry, were imaged using 488 nm and 561 nm wavelength lasers, respectively; membrane dye was imaged with 640 nm laser. Z-stacks were collected over the cell height with a step size allowing for oversampling. Multiple regions of interest were recorded within each well.

#### Point Scanning Confocal Microscopy

For characterization of FC puncta without PpIDs, super resolution microscopy was utilized with the Zeiss LSM 980 Airy Scan 2.0 microscope using a 100x α PlanApochromat 1.46 Oil DIC M27 objective. Imaging was carried out in the Airyscan SR mode via Zeiss Zen Blue 3.2 software. The mCherry-tagged FC and the membrane dye were imaged with 561 nm and 639 nm wavelength lasers. Z-stacks were collected over the cell height with a step size allowing for oversampling. Multiple regions of interest were recorded within each well.

### Image Analysis

#### Image Processing

Maximum projection Z-stacks for each channel were generated in ImageJ or Zeiss Zen Blue and used for all further analyses. Masks for cell segmentation were produced with CellPose based on the membrane dye signal. All images were processed with CellProfiler. Briefly, cell objects that were abnormally large or small were excluded. Furthermore, dead cells, as suggested by enriched membrane dye intake, were removed from analysis. [C]_T,Cell_ and [D]_T,Cell_ were approximated by the background subtracted average mCherry and GFP intensities, respectively, in each cellular mask. Pearson correlation coefficient was measured on a per cell basis within each cellular mask. Puncta masks were segmented with adaptive Sauvola thresholding of the mCherry signal. A dilute phase mask of each cell was taken to be the complement of the cell and puncta masks. [C]_T,Dilute_ and [D]_T,Dilute_ were approximated by the background subtracted average mCherry and GFP intensities, respectively, in each dilute phase mask. Dilute phase volume fraction was measured as the ratio of the number of pixels in the dilute phase mask to that of the whole cell. Pearson correlation coefficient and recruitment was calculated only in cells for which the dilute phase volume fraction was less than 1 (i.e. cells that had segmented puncta).

### Experimental Calculation of R

Recruitment was calculated experimentally as (see Supplemental Note 4.2):

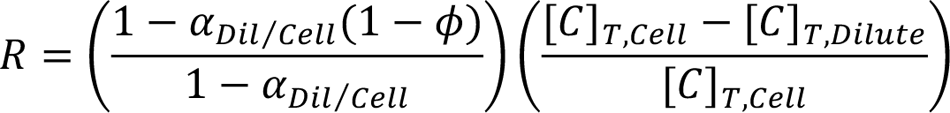

Where α*_Dil/Cell_* = [D]*_T,Dil_*⁄[D]*_T,Cell_* and (1 − ø) is the dilute phase volume fraction.

#### Calibration

yMW1099, expressing mCherry and GFP coupled by a self-cleaving 2A peptide from Equine rhinitis B virus^70^ was used to represent cells with equivalent concentrations of each fluorescent protein. Imaging yMW1099 was used to calibrate the relative intensities of mCherry and GFP at the imaging settings used.

#### Venus Protein-fragment Complementation Assay

The fragments of the Venus fluorescent protein were Venus_N_ (amino acids 1-210) and VenusC (amino acids 210-238)^55^. Venus_N_ was C-terminally tagged with PTB_P_, and Venus_C_ was N-terminally tagged with ALFA_P_. Triplicates of samples on three separate 24-well plates were outgrown as described for imaging sample preparation. At the 9-hour timepoint, fluorescent measurements and OD600 were collected with a monochromator-based plate reader (TECAN infinite200PRO MPlex). Selected excitation and emission Venus signal was 510nm/548nm. Fluorescent signal was normalized as previously described^71^ to account for background signal contributed by the media and cells. Three colonies of each strain were initially screened; the median performer was selected and tested in triplicate in three separate wells.

### Farnesene Fermentation

#### Strain Development

Starting with a mevalonate over-producing strain (JCY51)^72^, we utilized the Ellis lab CRISPR/Cas9 constructs^73^ to endogenously tag the ERG20 gene. The sgRNA sequence was designed to target the ERG20 coding region (5’-ATGTTCTTGGGTAATCAACA<u>AGG</u>). The repair template is consisted of a C-terminally PDZ_P_-tagged codon-optimized ERG20 from *S. cerevisiae* S288C and the upstream (583 bp) and downstream (500 bp) homologies of the *S. cerevisiae* CEN.PK2-1C ERG20. Colonies were screened using yeast colony PCR, and positive colonies were verified via PCR amplification and purification of the repaired region using primers outside of the homologies in the repair template, followed by sequencing, creating yKX489. To cure out the Cas9 plasmid, yKX489 was cultured overnight in YPD with 2% glucose and plated onto a fresh YPD plate. Colonies without the Cas9 plasmid were unable to grow in SC-knockout media, but were able to grow in SC-complete media, creating yKX485. To create a control strain without enzyme recruitment, two copies of FC* followed by two copies of SH3_P_-tagged aaFS_mut_ with two copies of FC* were sequentially integrated to the *his3* and *leu2* sites, respectively, creating yKX617. To create the basis for recruitment of the farnesene biosynthesis to engineered condensates, two copies of PDZ_D_-PDZ_D_-FC* followed by two copies of SH3_P_-tagged aaFS_mut_ with two copies of SH3_D_-SH3_D_-FC* were sequentially integrated to the *his3* and *leu2* sites, respectively, creating yKX604. From there, we used EasyClone2.0 vector pCFB2225 (a gift from Irina Borodina, Addgene plasmid # 67553) to incorporate additional FC* copies into the XII-2 locus (Chr XII: 808805..809939)^74^: 2 additional copies of SH3_D_-SH3_D_-FC* were integrated to the XII-2 site to make yKX646; 2 additional copies of PDZ_D_-PDZ_D_-FC* were integrated to the XII-2 site to make yKX647; 1 copy of SH3_D_-SH3_D_-FC* and 1 copy of PDZ_D_-PDZ_D_-FC* were integrated to the XII-2 site to make yKX656. In all cases, FC* uses I3-01(K129A) as the monomeric core unit in the FC construct with the FUS IDR and mCherry fluorescent protein^75^. All integrations are PCR verified.

#### Sample Preparation and Fermentation

Following yeast transformation, six colonies were randomly picked and verified for proper integration with PCR. Three positive isolates were grown overnight in 600 μL SC media lacking appropriate amino acid with 2% (w/v) glucose at 30°C in 24-well plate. On the next day, the overnight culture was used to inoculate 3 mL of SC media (Sunrise Science Products) with 2% (w/v) glucose in 14 mL test tube (Falcon) at a starting OD_600_ of 0.2 and incubated at 30°C for 144 hr with shaking at 265 rpm. 600 μL dodecane (Sigma Aldrich), was added to the culture after inoculation. Following fermentation analysis (as described below), the median performing colony was used for further strain development.

#### Analytical Methods and Analysis

To quantify farnesene production, 200 μL of the upper (dodecane) layer was sampled and diluted with 800 μL of ethyl acetate (Sigma Aldrich). The mixture was mixed for 5 min and centrifuged at 12,500 g for 10 min at 4°C to remove any residual aqueous phase and yeast cells. Then 500 μL of the supernatant was taken for GC-FID analysis.

β-farnesene was measured using an Agilent 7890B GC System. The concentration of the analyte was determined by comparing the peak areas with those of standard solutions. For analysis, samples were injected and subjected to a split (0.5 μL injection volume; 1:20 split) with a constant flow of 1.5 mL/min. Separation of samples was achieved using a DB-Wax column (30 m length, 0.25 mm diameter, 0.5 μm film) and a gradient method. Initially, the oven temperature was set to 70 °C and maintained for 5 minutes. Subsequently, the temperature was increased at a rate of 30 °C/min until reaching 230 °C, and then held for 3 minutes. Quantification of samples was performed using flame-ionization detection (300 °C, with H_2_ flow at 30 mL/min, air flow at 400 mL/min, and makeup flow at 25 mL/min), and the results were compared to a β-farnesene standard, a gift from Derek McPhee and Amyris.

#### Model

For the sake of modeling recruitment from the basis of equilibrium protein-ligand binding, R was rewritten in terms of the cellular concentration of bound cargo-domain assembly ([CD]_Cell_), as shown in Supplemental Note 4.3:

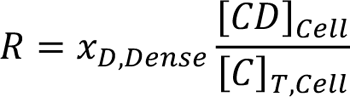

Given the assumptions of constant K_D_ and unbound cargo concentration, [CD]_Cell_ can be given directly by classical protein-ligand binding models^51^ (Supplemental Note 4.3).

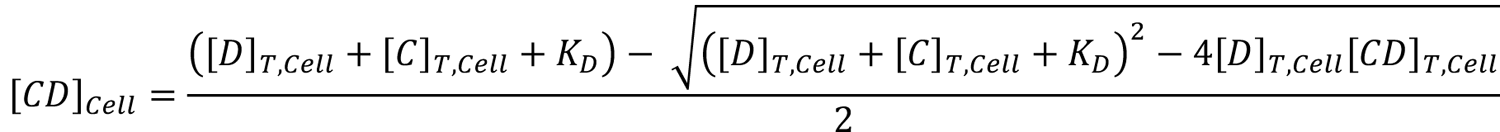

## Acknowledgements

We thank Evangelos Gatzogiannis for assistance with all microscopy experiments; Yi-Che Chang for advice on image analysis; Ushnish Rana for aiding in the development of the analytical model; Scott Wegner for assistance with strain development for and detection of β-farnesene; Derek McPhee and Amyris for providing β-farnesene standards; Kay Siu and Shannon Hoffman for thorough readings of the manuscript; and all members of the Avalos and Brangwynne labs for helpful advice, feedback, and discussions. This work was supported by the Howard Hughes Medical Institute, the Princeton Biomolecular Condensate Program, the AFOSR MURI (FA9550-20-1-0241), the U.S. Department of Energy, Office of Science, Office of Biological and Environmental Research (DE-SC0022155), and the Princeton Center for Complex Materials (PCCM), a U.S. National Science Foundation Materials Research Science and Engineering Center (Grant Nos. DMR-1420541 and DMR-2011750). M.T.W. is supported by the NSF GRFP (DGE-1656466).

## Author Contributions

M.T.W, C.P.B. and J.L.A conceptualized the PpID-FC system; M.T.W and K.X. performed experiments, which were analyzed by M.T.W.; strain development for β-farnesene production performed by K.X.; the manuscript was written by M.T.W. with contributions from all authors.

## Data and Code Availability Statement

Source data and code will be made available upon reasonable request to the corresponding authors.

## Extended Data for

**Extended Data Table.**
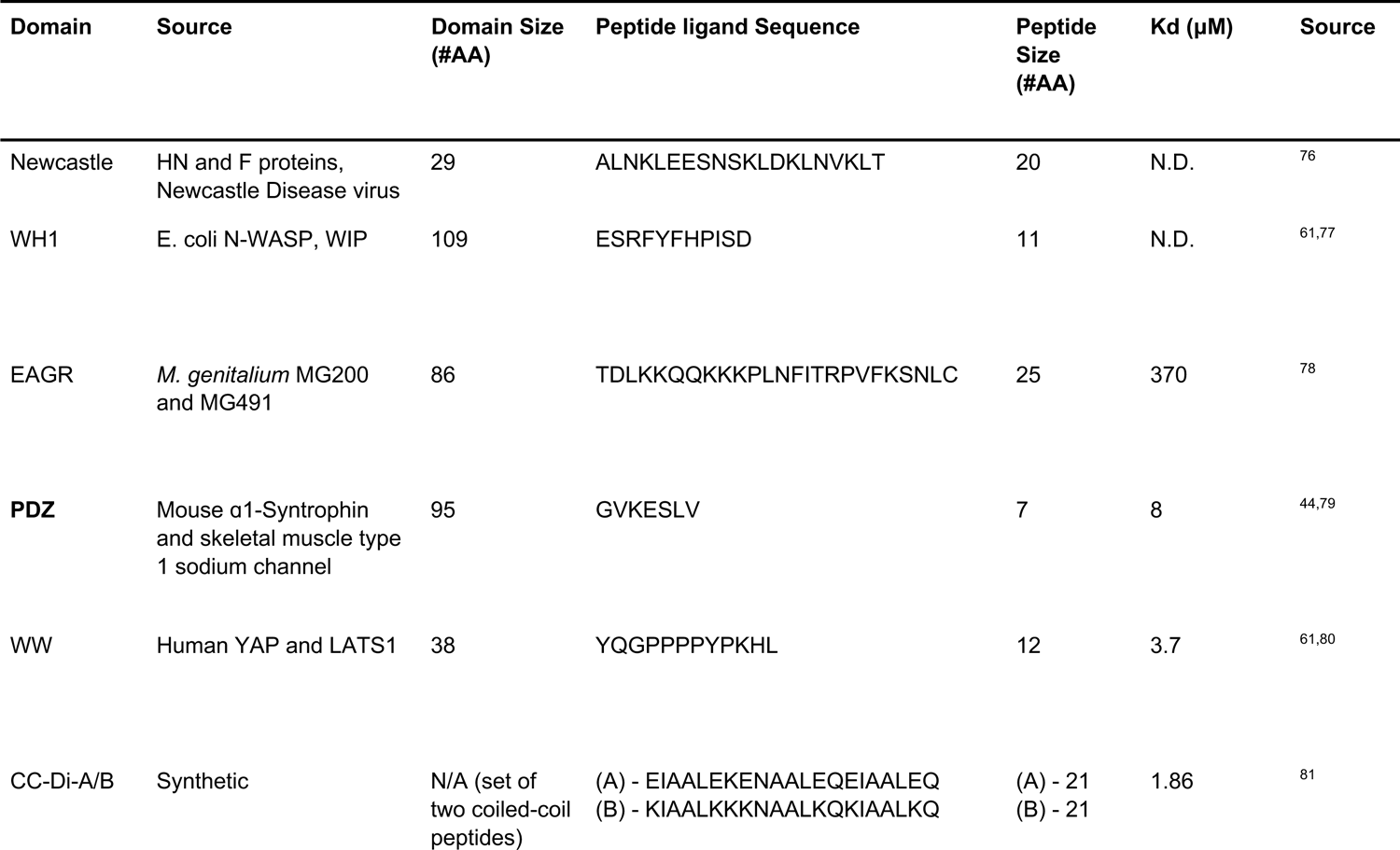
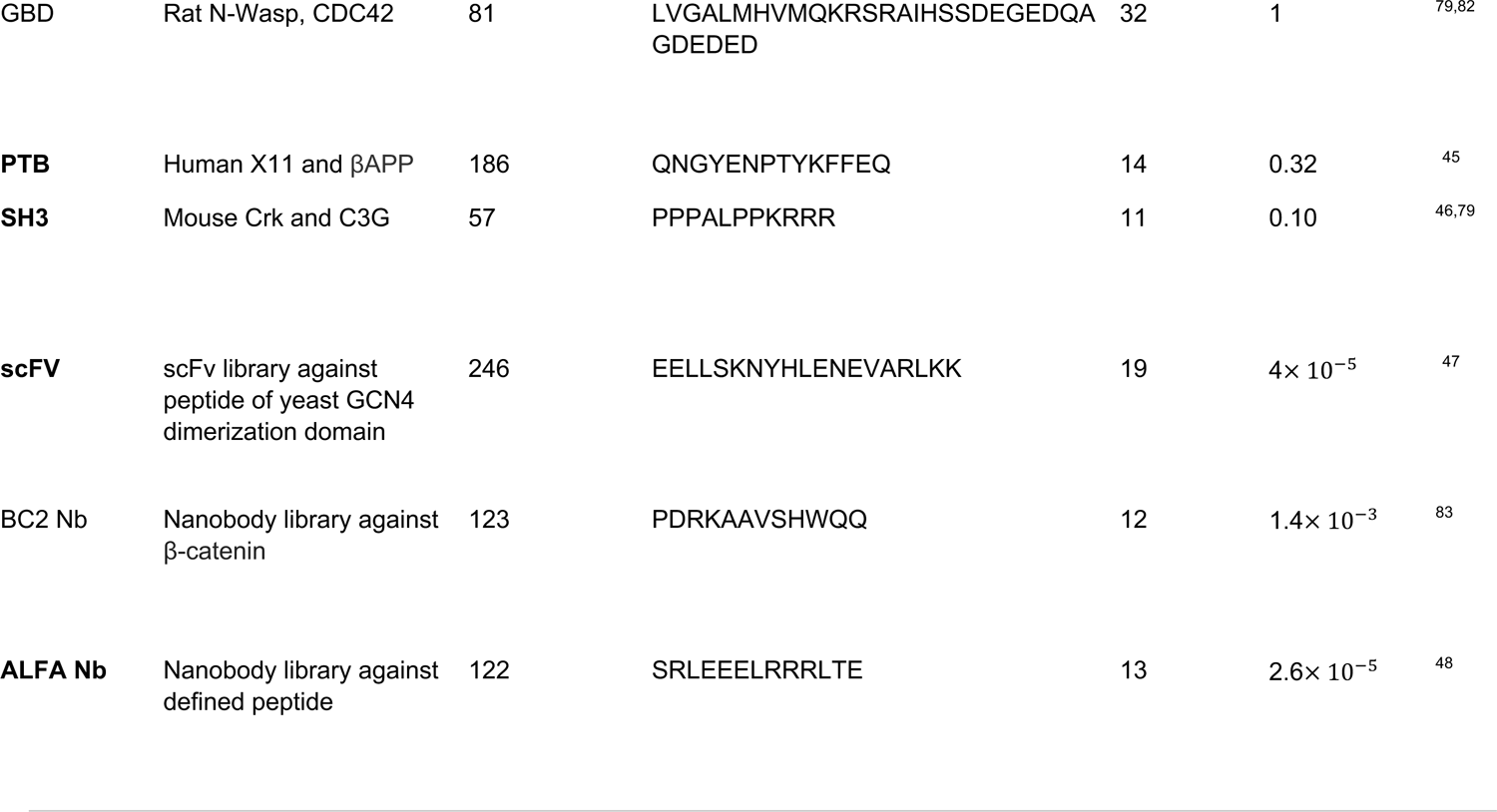
Characteristics of all tested PpIDs, listed in order of decreasing reported K_D_ N.D.: no data available. Bolded entries were utilized in this study.

**Extended Data Figure 1.**
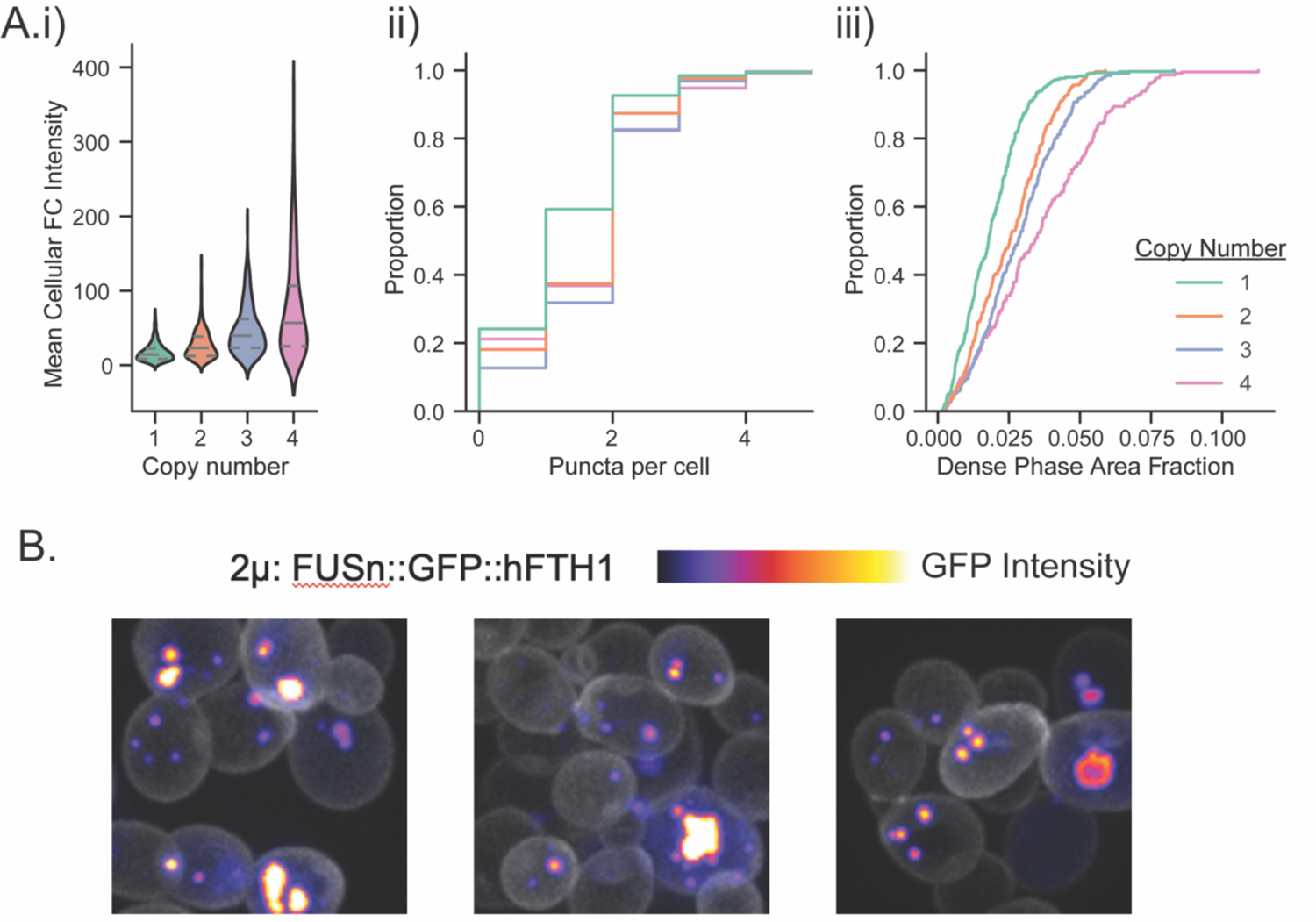
Additional FC Properties. A. i) Distribution of cell-averaged FC intensity for different expression conditions. Median, as well as first and third quartiles are marked as gray dashed lines; ii) Distribution of number of puncta in yeast expressing FC construct at multiple copy numbers; iii) Distribution of dense area fraction in yeast expressing FC construct at multiple copy numbers, not counting cells without detected puncta. (i-iii) For all, color scheme as indicated in (iii); n>280 cells for each strain B. Irregular puncta observed in 2μ (highly multi-copy and variable) expression of GFP-tagged FC

**Extended Data Figure 2.**
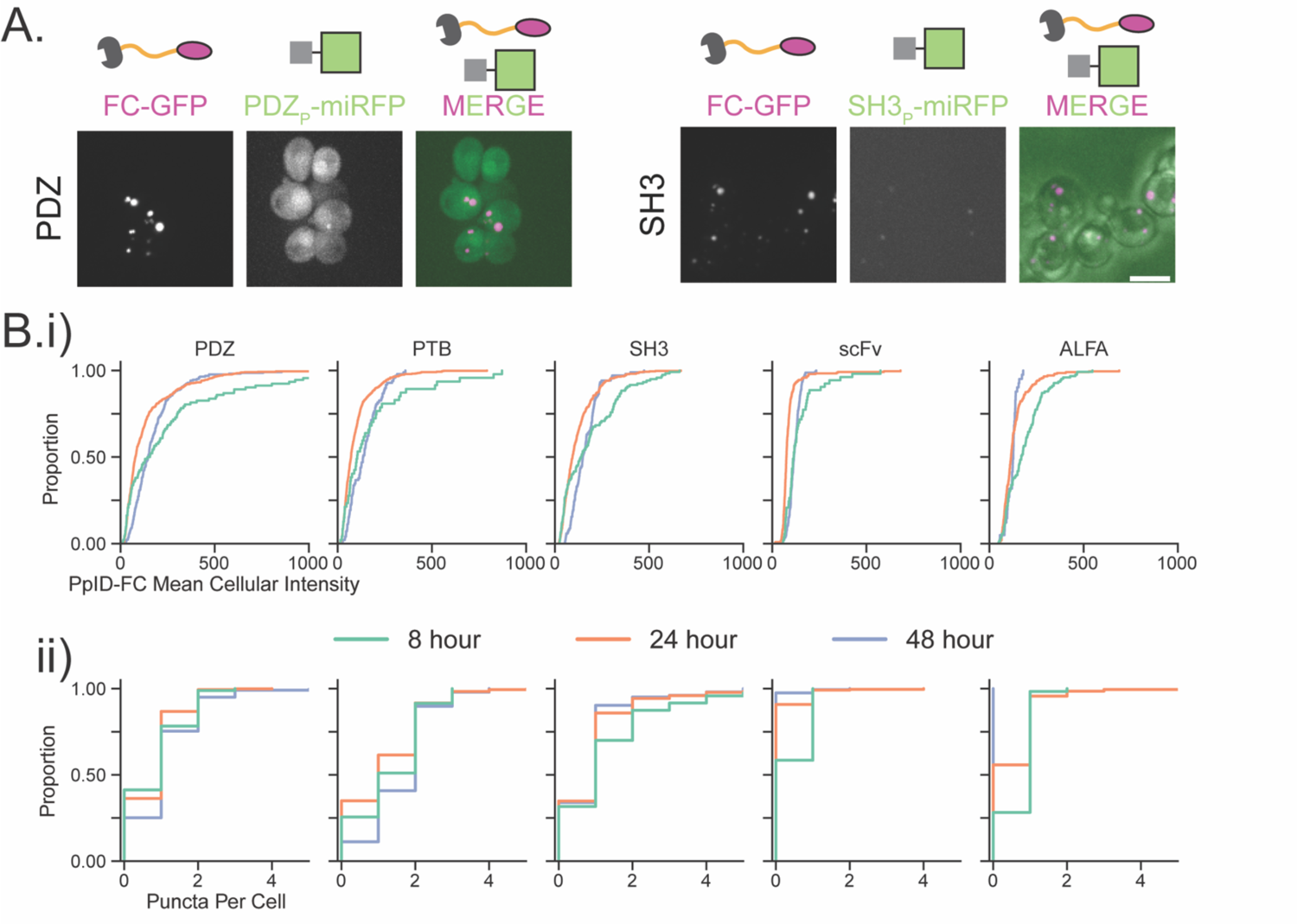
Additional PpID Properties. A. Representative images of initial PpID testing with 2µ expression of GFP-labelled FC with miRFP670 recruitment showing weak recruitment of N-terminally tagged PDZ_P_-miRFP and poor expression of SH3_P_-miRFP. DIC added in merged image for SH3 to show cell outlines. Scale bar, 5 µm B. Distributions of (mCherry tagged) PpID-FC expression, as assayed by (i) cellular mCherry intensity and (ii) cellular puncta count at different stages of growth. In all cases PpID-FC is coexpressed with GFP cargo C-terminally tagged with the corresponding peptide. Plots in (ii) correspond to the PpID label above it. scFv and ALFA PpIDs show notable decrease in expression and puncta count at later times. (n>40 cells for each strain and timepoint)

**Extended Data Figure 3.**
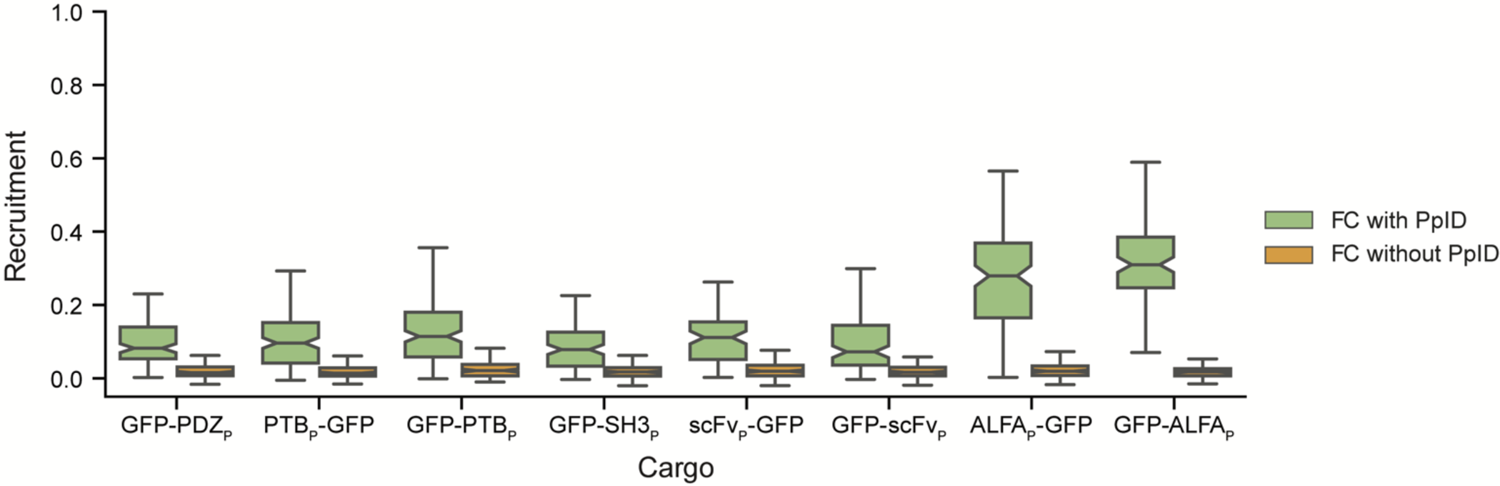
Population Recruitment for All PpID-peptide arrangements Distribution of recruitment for strains expressing GFP-peptide fusion with the corresponding PpID-FC (Green) or control without PpID fusion to FC (Orange). Arranged in order of decreasing reported K_D_ (See Extended Data Table for values). Boxes extend from the first to third quartiles, defining the interquartile range (IQR). Whiskers mark max and minimum value within 150% of the IQR from the box; outliers (exceeding 150% IQR from the box) are not shown. Notches show 95% confidence intervals of the median (center line). (n>80 cells for each strain)

**Extended Data Figure 4.**
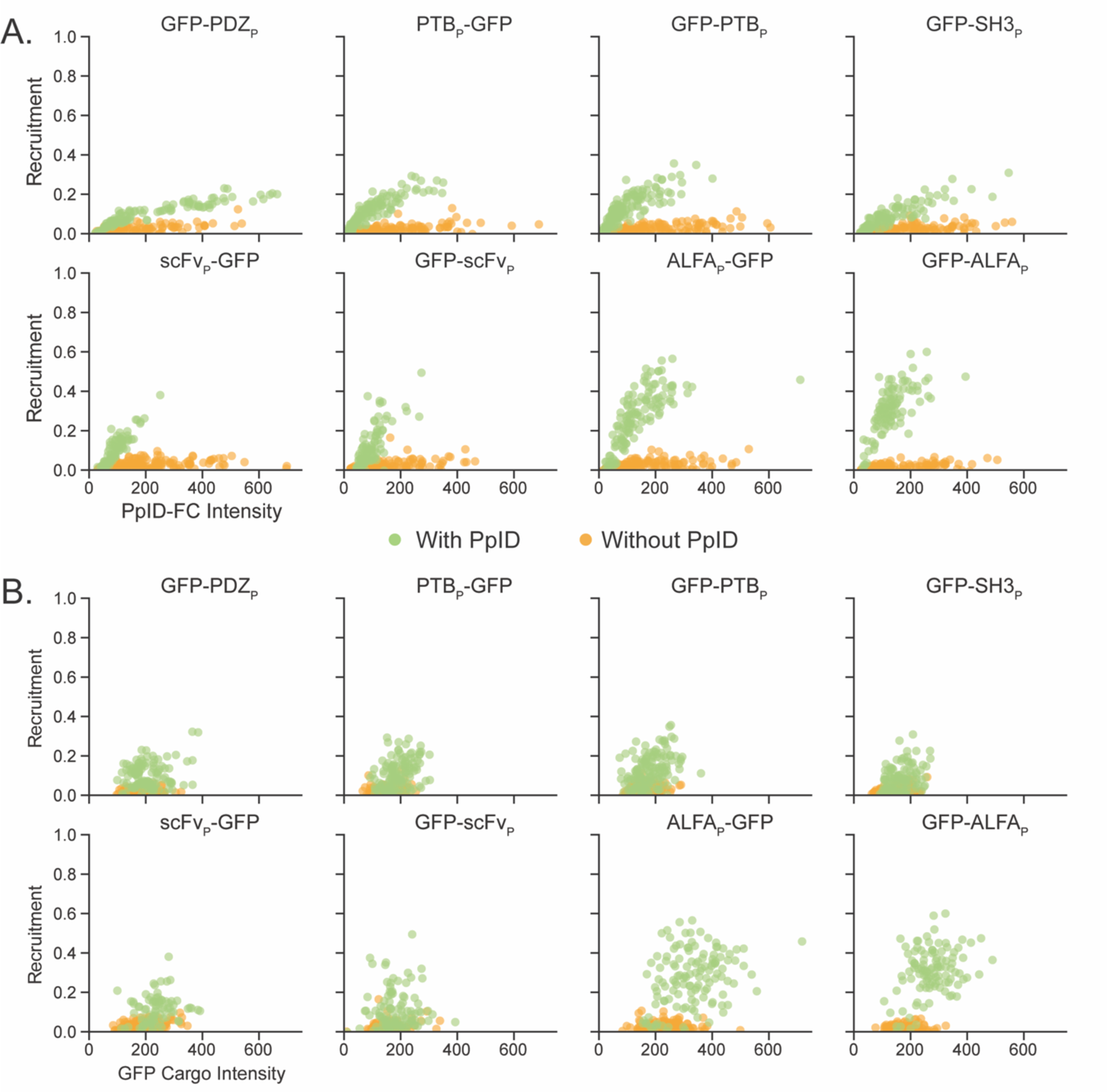
Recruitment dependence on PpID-FC and Cargo concentration for all PpIDs. A. Per-cell recruitment with respect to PpID-FC fluorescence Intensity for N and C terminal Cargo fusion. C-terminal fusion data replicated from Figure 3B(i) for ease of comparison B. Per-cell recruitment with respect to Cargo fluorescence Intensity for the same strains and datasets as in (A). Values of FC and Cargo intensities are normalized to account for the varying brightness of the fluorescent proteins used. (n>80 cells for each strain)

**Extended Data Figure 5.**
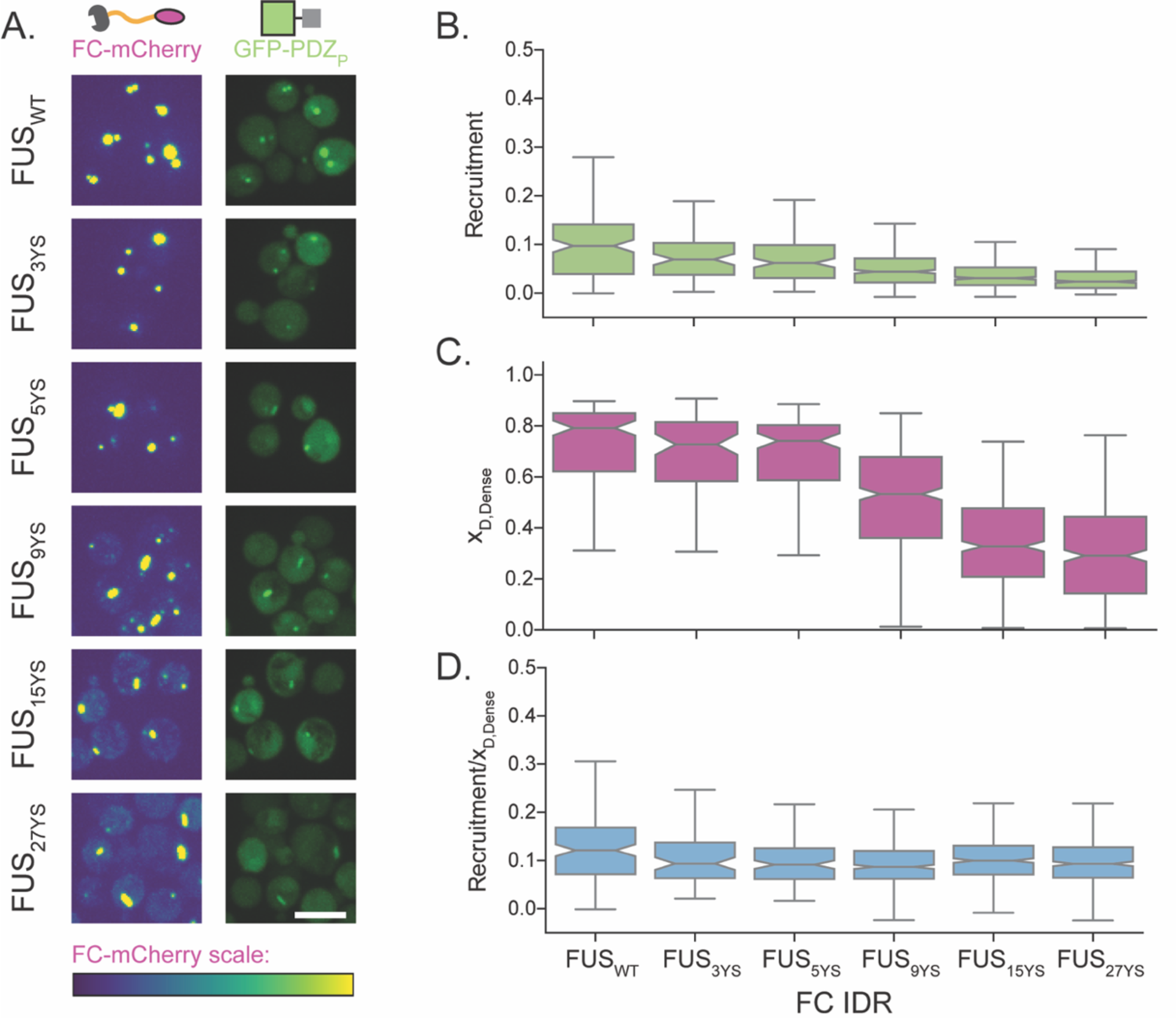
PDZ_D_-FC recruitment in clusters with decreasing x_D,Dense_. A. PDZ_D_-FC constructs where FUS IDR variants are used in the FC component with decreasing self-interaction through substitution of an increasing amount of tyrosine amino acids with serine, as indicated. Scale bar, 5 µm B. Recruitment of GFP-PDZ_P_ to the various PDZ_D_-FC constructs C. Increasing tyrosine to serine mutation leads to a decrease in x_D,Dense_ D. A conserved [CD]_Cell_/[C]_T,Cell_ ratio is observed by dividing the recruitment by x_D_,_Dense_ on a per cell basis. B-D) Labels as shown in D. Boxes extend from the first to third quartiles, defining the interquartile range (IQR). Whiskers mark max and minimum value within 150% of the IQR from the box; outliers (exceeding 150% IQR from the box) are not shown. Notches show 95% confidence intervals of the median (center line). (n>75 cells for each strain)

**Extended Data Figure 6.**
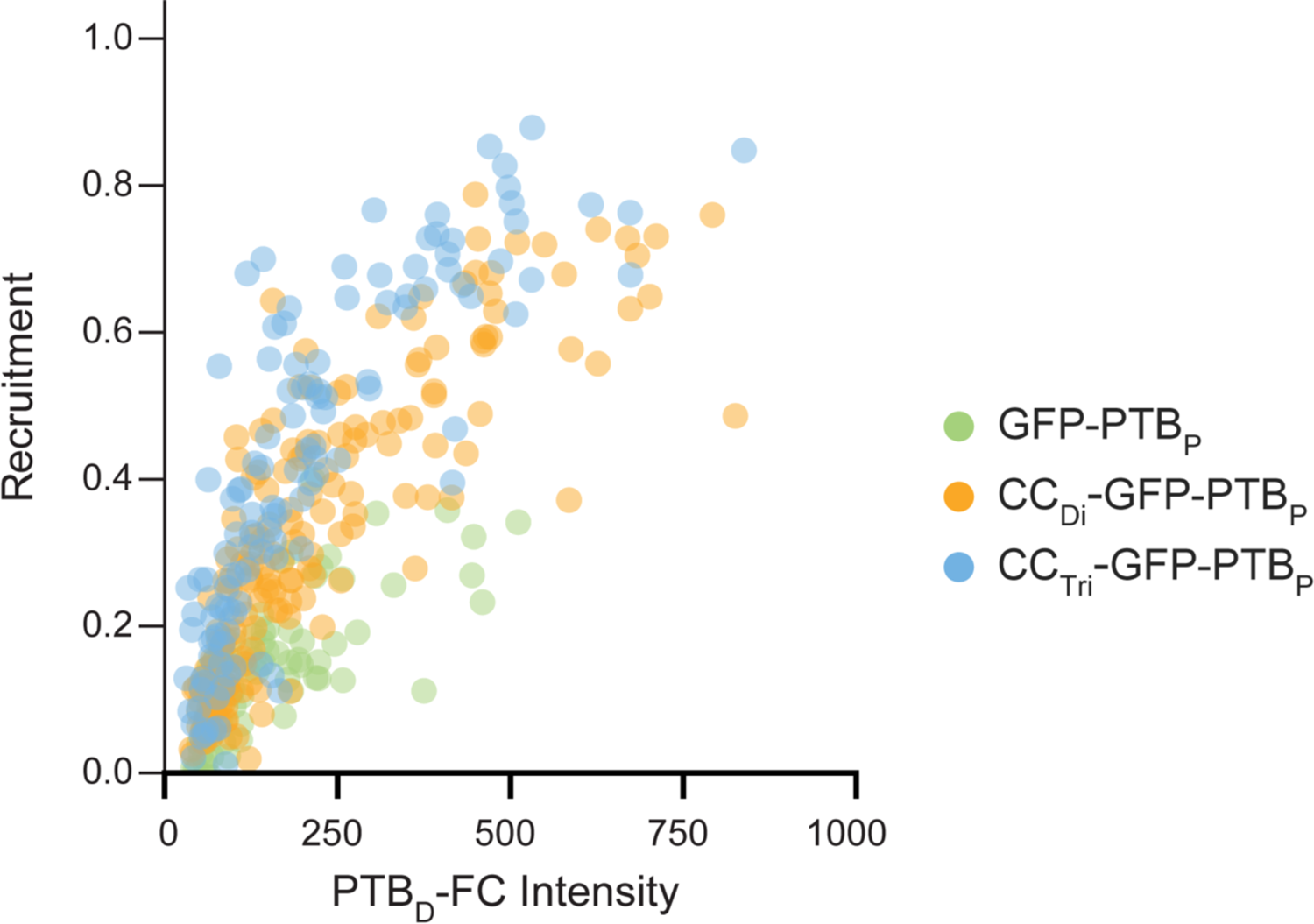
Cargo of increasing multimerization state showcase higher recruitment Cellular recruitment of GFP cargo to PTB_D_-FC for cargo of increasing multimerization. Multimeric proteins are mimicked with GFP-PTB_P_ cargo by adding engineered coiled coil domains to their N-terminus – Green: No N-terminal modification; Yellow: Dimeric coiled-coil; Blue: Trimeric coiled-coil. (n>70 cells for each strain)

**Extended Data Figure 7.**
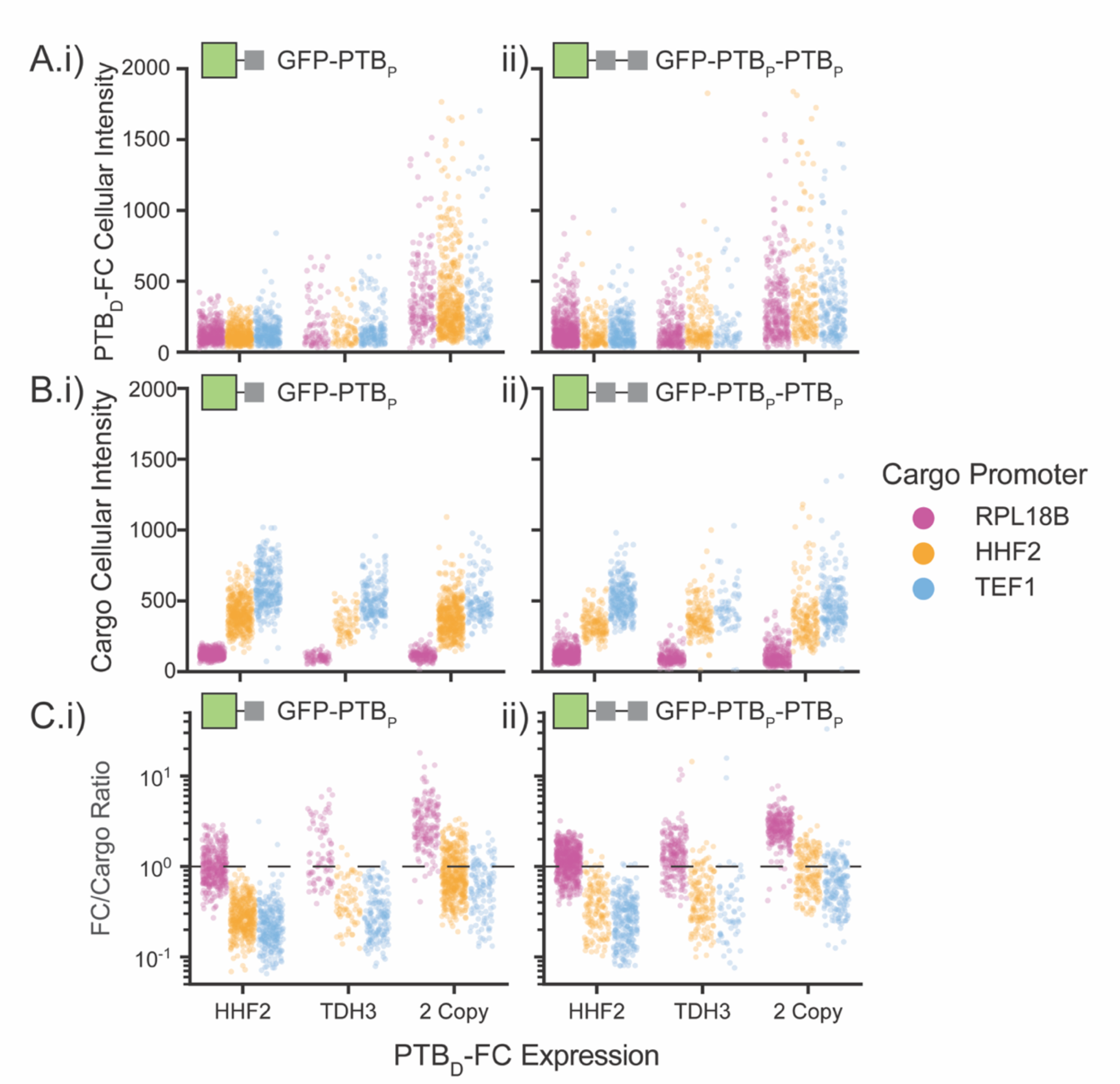
Cellular expression levels with PTB_D_-FC Recruitment Studies For the experiments shown in Fig 5, a range of expression levels were studied for PTB_D_-FC, and two forms of GFP cargo: _i_) GFP-PTB_P_ and ii) GFP-PTB_P_-PTB_P_. Three levels of expression of PTB_D_-FC were achieved by expression with the native HHF2 promoter, the native TDH3 promoter or with two copies, both driven by the TDH3 promoter. Three levels of expression of the cargo components were achieved using three native promoters of increasing strength: RPL18B, HHF2, or TEF1. These combinations resulted in the resulting expression distributions: A. Mean cellular intensity of PTB_D_-FC B. Mean cellular intensity of GFP Cargo C. PTB_D_-FC to GFP Cargo Cellular ratio Values of FC and Cargo intensities are normalized to account for the varying brightness of the fluorescent proteins used. (n>60 cells for each strain)

