## Supplemental Materials for "A Modular Design for Synthetic Membraneless Organelles Enables Compositional and Functional Control"

### Supplemental Notes

#### 1 Forever Corelet Characteristics

With increasing copy number of the FC construct, a broadened distribution of expression is observed (Extended Data Fig 1A(i)). Concurrently, the dense phase volume fraction shifts to higher values with a small change in the number of puncta (Extended Data Fig 1A(ii)) which leads to larger dense phase fraction with increasing FC expression (Extended Data Fig 1A(iii)). While Forever Corelets exhibit hallmarks of liquid-liquid phase separation (Fig 1C), at high levels of expression FC, material properties often appear more arrested and gel-like. Specifically, we observe irregular puncta when expressing FC when using the high copy number $2\mu$ plasmid (Extended Data Fig 1B). In sum, the FC condensates appear to assemble via liquid-liquid phase separation but are prone to exhibit gel-like properties, a feature common to many biomolecular condensates[1,2].

#### 2 PpID Selection

In keeping with the goal of using protein binders of small ligand tags, we searched for binders that interact with peptides of less than 35 residues (Extended Data Table). These included naturally occurring and engineered domains. Those that are naturally occurring largely originate from eukaryotic signaling cascades[3] and have been widely used in nonnative contexts to control protein localization[4], and exhibit reported dissociation constants, $K_D$, on the order of $10^{-1}$ to $10^{+1}$ $\mu$M. We also tested domains of engineered origin including a small chain variable fragment of an antibody, and nanobodies screened against small peptides with reported $K_D$ lower than $10^{-4}$ $\mu$M

#### 3 PpID Characteristics

PDZ$_P$ and SH3$_P$ only function as C-terminal fusions (Extended Data Fig 2A) owing to the PDZ$_D$ interaction involving the C-terminal -COOH[5], and the N-terminal expression of the proline-rich SH3$_P$ greatly diminishing cargo expression. The ALFA$_D$ and scFv$_D$ domains show instability in older cells,i.e. cells in later stage of growth, in contrast, PDZ$_D$, SH3$_D$, and PTB$_D$, exhibit largely unchanged behavior, well into the stationary phase (Extended Data Fig 2B).

### 4 Recruitment Derivations

#### 4.1 Definitions

For cargo ($C$) and FC-PpID concentration ($D$), the following is true within any cellular region i, where i can be the condensate phase ($Dense$), dilute phase ($Dil$), or whole cell system ($Cell$):

$$D_{T,i} = D_{F,i} + [CD]_i \tag{S-1}$$

$$C_{T,i} = C_F + [CD]_i \tag{S-2}$$

Where subscript $T$ refers to Total, or sum, of the unbound, free population (subscript $F$) and bound population of cargo and FC-PpID ($CD$). While all terms are concentration terms, square bracket notation is used only in the case of the bound fraction, for clarity. Note that $C_F$ requires no region label as it is assumed to be equivalent in all portions of the cell.

As a result of mass balance,

$$X_{Cell} = \phi X_{Dense} + (1 - \phi)X_{Dil} \tag{S-3}$$

where $X$ can be $D_T$,$D_F$,$[CD]$,$C_T$, or $C_F$ and $\phi$ represents the dense volume fraction.

Equilibrium $D$-$C$ interaction (as mediated directly through domain-peptide interaction) is expressed in terms of the equilibrium dissociation constant, $K_D$, which is assumed to be equivalent over all regions. We can therefore write

$$K_D = \frac{D_{F,Dense}C_F}{[CD]_{Dense}} = \frac{D_{F,Dil}C_F}{[CD]_{Dil}} = \frac{D_{F,Cell}C_F}{[CD]_{Cell}} \tag{S-4}$$

#### 4.2 Definition of recruitment for experimental measurements

We are interested in measuring the bound portion of cargo in the dense phase as this is a direct indication of the engineered degree of protein enrichment. We therefore define recruitment ($R$) as the fraction of bound cargo molecules in the dense phase (from Equation 1 in the main text):

$$R = \frac{N_{CD,Dense}}{N_{C_T,Cell}} = \phi\frac{[CD]_{Dense}}{C_{T,Cell}} \tag{S-5}$$

where $N_{CD,Dense}$ represents the number of cargo molecules bound to domain molecules in the dense phase and $N_{C_T,Cell}$ represents the total number of cargo molecules (regardless of bound state) in the entire cell.

Experimentally, we are only able to measure 'Total' concentrations of domain ($D_T$) and cargo ($C_T$) in the dilute regions of the cell, as well as the entirety of the cell. Measurement in the dense phase is more challenging in some cases when puncta are small and similar in size to the diffraction limit. For this reason, we focus only on concentrations, as inferred by mean fluorescence intensity, in the whole cell and dilute cellular region and measurements of dilute phase volume fraction, $(1 - \phi)$.

Given the assumption that $C_F$ is constant throughout the cell, we can remove its contribution to the cargo concentrations by writing from Equation S-2:

$$C_{T,Cell} - C_{T,Dilute} = [CD]_{Cell} - [CD]_{Dilute} \tag{S-6}$$

Given assumptions that the equilibrium binding ($K_D$) and unbound cargo concentration ($C_F$) are equivalent in all regions, from Equations S-1 and S-4, $[CD]_{Dilute}$ can be written in terms of $[CD]_{Cell}$:

$$\frac{D_{T,Cell} - [CD]_{Cell}}{[CD]_{Cell}} = \frac{D_{T,Dil} - [CD]_{Dil}}{[CD]_{Dil}}$$

$$[CD]_{Dil} = \underbrace{\frac{D_{T,Dil}}{D_{T,Cell}}}_{\alpha_{Dil/Cell}}[CD]_{Cell} \tag{S-7}$$

We are therefore able to directly infer the cellular bound population $[CD]_{Cell}$ in terms of experimentally measurable 'Total' concentrations by combining Equations S-6 and S-7 :

$$[CD]_{Cell} = \frac{C_{T,Cell} - C_{T,Dilute}}{1 - \alpha_{Dil/Cell}} \tag{S-8}$$

The recruitment metric (Equation S-5) can be recast by mass balance on [CD] (Equation S-3):

$$R = \frac{[CD]_{Cell} - (1-\phi)[CD]_{Dilute}}{C_{T,Cell}} = (1 - \alpha_{Dil/Cell}(1-\phi))\frac{[CD]_{Cell}}{C_{T,Cell}} \tag{S-9}$$

allowing for a final calculation of recruitment ($R$) (the fraction of bound cargo molecules in the dense phase) as

$$R = \left(\frac{1 - \alpha_{Dil/Cell}(1-\phi)}{1 - \alpha_{Dil/Cell}}\right)\left(\frac{C_{T,Cell} - C_{T,Dilute}}{C_{T,Cell}}\right) \tag{S-10}$$

All of these values are reliably measured experimentally through fluorescence microscopy. Therefore, Equation S-10 provides the means for all experimental measurements of recruitment reported in this work.

#### 4.3 Recruitment in terms of Cargo-Domain Binding Equilibrium

We intend to develop an analytical framework to predict recruitment as a function of (1) cellular concentrations of cargo and domain, (2) domain partitioning between dense and dilute phases of defined volumetric fraction, and (3) binding affinity. An equilibrium ligand-receptor binding framework can be used to determine the concentration of the cargo-domain bound fraction (CD). This is most readily applied with the expression of recruitment in terms of cellular concentrations. From Equation S-9

$$R = \underbrace{(1 - \alpha_{Dil/Cell}(1-\phi))}_{x_{D,Dense} = \phi\frac{[D]_{T,Dense}}{[D]_{T,Cell}}}\frac{[CD]_{Cell}}{C_{T,Cell}} \tag{S-11}$$

which is equivalent to Equation 2 in the main text. $x_{D,Dense}$ is the ratio of all domain molecules in the dense phase to all domain molecules in the cell.

To determine $[CD]_{Cell}$, we combine equations S-1, S-2, and S-4, and write

$$K_D = \frac{(D_{T,Cell} - [CD]_{Cell})(C_{T,Cell} - [CD]_{Cell})}{[CD]_{Cell}} \tag{S-12}$$

which leads to a quadratic equation:

$$[CD]_{Cell}^2 - (D_{T,Cell} + C_{T,Cell} + K_D)[CD]_{Cell} + D_{T,Cell}C_{T,Cell} = 0 \tag{S-13}$$

where only one solution is physically meaningful:

$$[CD]_{Cell} = \frac{(D_{T,Cell} + C_{T,Cell} + K_D) - \sqrt{(D_{T,Cell} + C_{T,Cell} + K_D)^2 - 4D_{T,Cell}C_{T,Cell}}}{2} \tag{S-14}$$

Equation S-14 combined with equation S-11 provides a full model for the fraction of bound cargo molecules in the dense phase as a function of whole cell concentrations of cargo and domain, the equilibrium dissociation constant, and the dense phase volume fraction:

$$R = x_{D,Dense}\frac{(D_{T,Cell} + C_{T,Cell} + K_D) - \sqrt{(D_{T,Cell} + C_{T,Cell} + K_D)^2 - 4D_{T,Cell}C_{T,Cell}}}{2C_{T,Cell}} \tag{S-15}$$

We can evaluate the partial derivative of R as a function of cellular domain concentration ($D_{T,Cell}$), under the assumption of constant $x_{D,Dense}$:

$$\left.\frac{\partial R}{\partial D_{T,Cell}}\right|_{x_{D,Dense},const.} = x_{D,Dense}\left[\frac{(C_{T,Cell} - D_{T,Cell} - K_D) + \sqrt{(D_{T,Cell} + C_{T,Cell} + K_D)^2 - 4D_{T,Cell}C_{T,Cell}}}{2C_{T,Cell}\sqrt{(D_{T,Cell} + C_{T,Cell} + K_D)^2 - 4D_{T,Cell}C_{T,Cell}}}\right] \tag{S-16}$$

which when evaluated at $D_{T,Cell} = 0$ gives

$$\left.\frac{\partial R}{\partial D_{T,Cell}}\right|_{x_{D,Dense}const.,D_{T,Cell}=0} = x_{D,Dense}\left[\frac{1}{C_{T,Cell} + K_D}\right] \tag{S-17}$$

which is Equation 3 in the main text.

#### 4.4 Recruitment behavior in limiting regimes of $[C]_{T,Cell}/K_D$

If we consider $[CD]_{cell} \ll \sqrt{D_{T,Cell}C_{T,Cell}}$ (which is largely true outside of the region of $C_{T,Cell}/D_{T,Cell} = 1$ and $K_D/D_{T,Cell} \ll 1$(see Figure S1)), then we can consider the quadratic term in Equation S-15 to be negligible and approximate R as:

$$\frac{R}{x_{D,Dense}} = \frac{D_{T,Cell}}{D_{T,Cell} + C_{T,Cell} + K_D} \tag{S-18}$$

In the case of $C_{T,Cell} \ll D_{T,Cell} + K_D$, we further approximate R as

$$\frac{R}{x_{D,Dense}} = \frac{D_{T,Cell}}{D_{T,Cell} + K_D} \tag{S-19}$$

which is Equation 4 in the main text.

In the case of $D_{T,Cell} + K_D \ll C_{T,Cell}$, we further approximate R as

$$\frac{R}{x_{D,Dense}} = \begin{cases} \frac{D_{T,Cell}}{C_{T,Cell}} & \text{if } D_{T,Cell} < C_{T,Cell} \\ 1 & \text{else} \end{cases} \tag{S-20}$$

which is Equation 5 in the main text. The requirement of $D_{T,Cell} < C_{T,Cell}$ is added as otherwise Equation S-20 would give values of $R > x_{D,Dense}$, which is not physically meaningful.

In the main text we neglect the presence of $D_{T,Cell}$ in the inequalities that dictate the applicable regime approximation (i.e. $C_{T,Cell} \ll$ or $\gg K_D$ instead of $C_{T,Cell} \ll$ or $\gg \mathbf{D_{T,Cell}} + K_D$) to make evident the interplay between $C_{T,Cell}$ and $K_D$ when $D_{T,Cell}$ is considered as the independent variable and considered over a large range of values. This omission does not alter the interpretation discussed in the main text.

In the case of $C_{T,Cell} \ll D_{T,Cell} + K_D$, neglecting $D_{T,Cell}$ under-approximates the regime in which Equation S-19 is applicable, as suggested by the fact that Equation S-19 yields a reasonable approximation of R for $C_{T,Cell}/K_D = 1$, as shown in main text Figure 3D. In the case of $D_{T,Cell} + K_D \ll C_{T,Cell}$, Equation S-20 is only valid for $D_{T,Cell} < C_{T,Cell}$; therefore it is most important to consider the relative values of $K_D$ and $C_{T,Cell}$ to determine if Equation S-20 will provide a reasonable approximation in the regime of $D_{T,Cell} < C_{T,Cell}$.

### 5 Venus Protein-fragment Complementation Assay

The Venus PCA used in this study fragments Venus at amino acid 210 (ref. 6), resulting a 210 amino acid fragment (Venus$_N$), C-terminally tagged with PTB$_P$, and a 29 amino acid fragment (Venus$_C$), N-terminally tagged with ALFA$_P$. Our PCA experiment does show sensitivity of the bulk Venus signal on whether the ALFA$_P$-Venus$_C$ fragment is recruited to FC condensates (see 3rd and 4th bars of Fig 4C). We hypothesize that the small Venus$_C$ fragment may experience a reduced half-life due to instability, that could be increased upon recruitment with partial protection from the cellular degradation machinery. Therefore, to avoid confounding interpretations, in testing stoichiometric control, we only consider changing the degree of recruitment of Venus$_N$-PTB$_P$ via varying expression of PTB$_D$-FC, while keeping ALFA$_D$-FC highly expressed (Fig4C, last 2 bars)

### 6 FC Construct Used for Farnesene Application

Showcasing the flexibility of the FC clustering system, an alternative 'Core' unit was used to make condensates for the recruitment of aaFS$_{mut}$ and ERG20. Initial experiments showed sensitivity of the production of some terpenes to ferritin production, potentially as a result of perturbing the level of iron homeostasis. To avoid this affect, the FTH1 in the FC construct is substituted with the I3-01(K129A) monomer, 60 of which self-assemble to form a dodecahedral protein assembly[7]. We have previously observed this substitution to form puncta similar to FTH1-based FC in yeast[8]. To accentuate the distinct design used here, we refer to the I3-01-based FC (FUSn::mCherry::I3-01(K129A)) as FC*. To further enhance recruitment, each FC* is N-terminally tagged with a tandem repeat of two PpIDs (either PDZ$_D$ or SH3$_D$), further increasing domain concentration, and therefore recruitment.

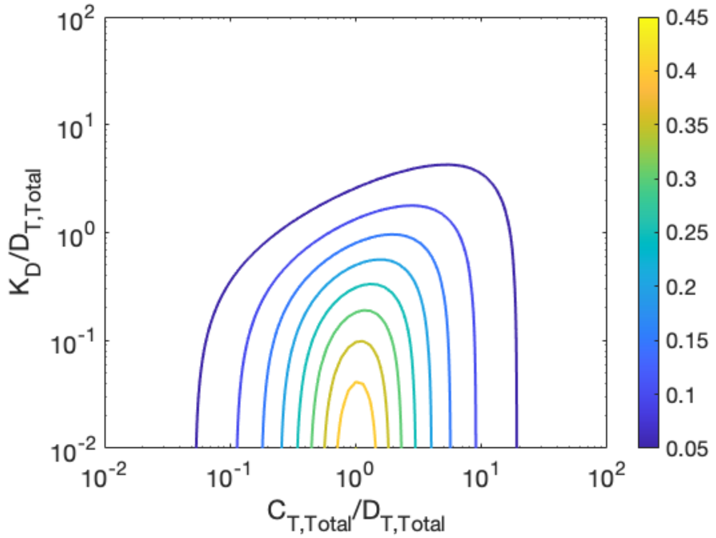


Figure S1: Contour plot of $\frac{[CD]_{Cell}}{\sqrt{D_{T,Cell}C_{T,Cell}}}$ where $[CD]_{Cell}$ is given by equation S-14

**Supplemental Table 1.** Plasmids Used in this study

| PLASMID | CODING SEQUENCE | MARKER | VECTOR | SOURCE |
|---|---|---|---|---|
| pKX005 | pHHF2_ALFAP-meGFP_tACT1 | LEU2 | Leu2 Integration | This Study |
| pKX006 | pHHF2_meGFP-ALFAP_tACT1 | LEU2 | Leu2 Integration | This Study |
| pKX038 | pTEF1_PDZD-FUSn-mCherry-FTH1_tTDH1 | HIS3 | His3 Integration | This Study |
| pKX039 | pTEF1_ALFAD-FUSn-mCherry-FTH1_tTDH1 | HIS3 | His3 Integration | This Study |
| pKX044 | pTEF1_SH3D-FUSn-mCherry-FTH1_tTDH1 | HIS3 | His3 Integration | This Study |
| pKX045 | pTEF1_PTBD-FUSn-mCherry-FTH1_tTDH1 | HIS3 | His3 Integration | This Study |
| pKX046 | pTEF1_scFvD-FUSn-mCherry-FTH1_tTDH1 | HIS3 | His3 Integration | This Study |
| pKX048 | pHHF2_PTBP-meGFP_tACT1 | LEU2 | Leu2 Integration | This Study |
| pKX049 | pHHF2_meGFP-PTBP_tACT1 | LEU2 | Leu2 Integration | This Study |
| pKX050 | pHHF2_scFvP-meGFP_tACT1 | LEU2 | Leu2 Integration | This Study |
| pKX051 | pHHF2_meGFP-scFvP_tACT1 | LEU2 | Leu2 Integration | This Study |
| pKX052 | pHHF2_meGFP-SH3P_tACT1 | LEU2 | Leu2 Integration | This Study |
| pKX053 | pHHF2_meGFP-PDZP_tACT1 | LEU2 | Leu2 Integration | This Study |
| pKX100 | pTEF1_FUSn-mCherry-FTH1_tTDH1 | HIS3 | His3 Integration | This Study |
| pKX323 | pTDH3_PDZD-PDZD-FUSn-mCherry-I301(K129A)_tTDH1, pTEF1_PDZD-PDZD-FUSn-mCherry-I301(K129A)_tTDH1 | HIS3 | His3 Integration | This Study |
| pKX325 | pTDH3_FUSn-mCherry-I301(K129A)_tADH1t, pTEF1p_FUSn-mCherry-I301(K129A)_tTDH1 | HIS3 | His3 Integration | This Study |
| pKX352 | pTEF1_aaFSmut-Myc-SH3P_tSSA1, pTDH3_SH3D-SH3D-FUSn-mCherry-I301(K129A)_tTDH1, pTDH3_aaFSmut-Myc-SH3P_tACT1, pCCW12_SH3D-SH3D-FUSn-mCherry-I301(K129A)_tENO1 | LEU2 | Leu2 Integration | This Study |

| | | | | |
|---|---|---|---|---|
| pKX382 | pTEF1_aaFSmut-Myc-SH3P_tSSA1, pTDH3_FUSn-mCherry-I301(K129A)_tTDH1, pTDH3_aaFSmut-Myc-SH3P_tACT1, pCCW12_FUSn-mCherry-I301(K129A)_tENO1 | LEU2 | Leu2 Integration | This Study |
| pKX386 | pTDH3_SH3D-SH3D-FUSn-mCherry-I301(K129A)_tTDH1, pCCW12_SH3D-SH3D-FUSn-mCherry-I301(K129A)_tENO1 | URA3 | XII-2 Integration | This Study |
| pKX517 | pCCW12_PDZD-PDZD-FUSn-mCherry-I301(K129A)_tENO1 pTDH3_PDZD-PDZD-FUSn-mCherry-I301(K129A)_tTDH1 | URA3 | XII-2 Integration | This Study |
| pKX557 | ERG20WT sgRNA | NA | CRISPR Guide | This Study |
| pKX532 | ERG20-Myc-PDZP | N/A | CRISPR Repair | This Study |
| pKX563 | pCCW12_PDZD-PDZD-FUSn-mCherry-I301(K129A)_ENO1t pTDH3_SH3D-SH3D-FUSn-mCherry-I301(K129A)_tTDH1 | URA3 | XII-2 Integration | This Study |
| pMW042 | pTEF1_FUSn-meGFP-FTH1_tACT1 | URA3 | 2µ Plasmid | This Study |
| pMW091 | pTDH3_SH3D-FUSn-meGFP-FTH1_tADH1 | URA3 | 2µ Plasmid | This Study |
| pMW092 | pTDH3_PDZD-FUSn-meGFP-FTH1_tADH1 | URA3 | 2µ Plasmid | This Study |
| pMW517 | pHHF2_meGFP_tACT1 | LEU2 | Integration | This Study |
| pMW699 | pTDH3_PDZD-FUSn-mCherry-FTH1_tTDH1 | HIS3 | His3 Integration | This Study |
| pMW728 | pTEF1_FUSn-mCherry-FTH1_tACT1 | URA3 | CEN/ARS Plasmid | This Study |
| pMW732 | pTDH3_PTBD-FUSn-mCherry-FTH1_tTDH1 | HIS3 | His3 Integration | This Study |
| pMW749 | pTEF1_FUSn-mCherry-FTH1_tTDH1 | LEU2 | Leu2 Integration | This Study |
| pMW750 | pTEF1_FUSn-mCherry-FTH1_tTDH1 | TRP1 | Trp1 Integration | This Study |
| pMW777 | pRPL18B_meGFP-PTBP_tACT1 | LEU2 | Leu2 Integration | This Study |
| pMW779 | pHHF2_meGFP-PTBP_tACT1 | LEU2 | Leu2 Integration | This Study |
| pMW781 | pTEF1_meGFP-PTBP_tACT1 | LEU2 | Leu2 Integration | This Study |

| | | | | |
|---|---|---|---|---|
| pMW788 | pCCW12_ALFAD-FUSn-mCherry-FTH1_tENO1, pTDH3_PTBD-FUSn-mCherry-FTH1_tTDH1 | HIS3 | His3 Integration | This Study |
| pMW789 | pHHF2_PTBD-FUSn-mCherry-FTH1_tTDH1 | HIS3 | His3 Integration | This Study |
| pMW837 | pTDH3_PTBD-FUSn-mCherry-FTH1_tENO1, pTDH3_PTBD-FUSn-mCherry-FTH1_tTDH1 | HIS3 | His3 Integration | This Study |
| pMW840 | pTDH3_FUSn-mCherry-FTH1_tTDH1, pCCW12_FUSn-mCherry-FTH1_tENO1 | HIS3 | His3 Integration | This Study |
| pMW845 | pHHF2_ALFAP-Myc-VenusC_tACT1, pHHF2_VenusN-HA-PTBP_tADH1 | LEU2 | Leu2 Integration | This Study |
| pMW888 | pCCW12_ALFAD-FUSn-mCherry-FTH1_tENO1, pHHF2_PTBD-FUSn-mCherry-FTH1_tTDH1 | HIS3 | His3 Integration | This Study |
| pMW902 | pTDH3_PDZD-FUSn(3YS)-mCherry-FTH1_tTDH1 | HIS3 | His3 Integration | This Study |
| pMW903 | pTDH3_PDZD-FUSn(5YS)-mCherry-FTH1_tTDH1 | HIS3 | His3 Integration | This Study |
| pMW904 | pTDH3_PDZD-FUSn(9YS)-mCherry-FTH1_tTDH1 | HIS3 | His3 Integration | This Study |
| pMW905 | pTDH3_PDZD-FUSn(15YS)-mCherry-FTH1_tTDH1 | HIS3 | His3 Integration | This Study |
| pMW906 | pTDH3_PDZD-FUSn(27YS)-mCherry-FTH1_tTDH1 | HIS3 | His3 Integration | This Study |
| pMW920 | pCCW12_ALFAD-FUSn-mCherry-FTH1_tENO1, pTDH3_FUSn-mCherry-FTH1_tTDH1t | HIS3 | His3 Integration | This Study |
| pMW921 | pCCW12_FUSn-mCherry-FTH1_tENO1, pTDH3_PTBD-FUSn-mCherry-FTH1_tTDH1 | HIS3 | His3 Integration | This Study |
| pMW989 | pHHF2_BCRDimer-meGFP-PTBP_tACT1 | LEU2 | Leu2 Integration | This Study |
| pMW992 | pHHF2_CCTrimer-meGFP-PTBP_tACT1 | LEU2 | Leu2 Integration | This Study |
| pMW998 | pRPL18B_meGFP-PTBP-PTBP_tACT1 | LEU2 | Leu2 Integration | This Study |
| pMW999 | pHHF2_meGFP-PTBP-PTBP_tACT1 | LEU2 | Leu2 Integration | This Study |

| | | | | |
|---|---|---|---|---|
| pMW1000 | pTEF1_meGFP-PTBP-PTBP_tACT1 | LEU2 | Leu2 Integration | This Study |
| pMW1003 | pTDH3_mCherry-(ERBV-1)2A-GFP_tTDH1 | HIS3 | His3 Integration | This Study |
| pYZ163 | EMPTY | HIS3 | His3 Integration | This Study |

**Supplemental Table 2**. Yeast strains used in this study

| YEAST STRAIN | GENOTYPE | FIGURES | SOURCE |
|---|---|---|---|
| CEN.PK2-1C | MATa his3Δ1 leu2-3_112 trp1-289 ura3-53 | | [9] |
| JCY51 | CEN.PK2-1C, YARCdelta5-(pTDH3-EfMvaE-tADH1, pTEF1-EfMvaS-tACT1, pPGK1-SeAcs(L641P)-tCYC1) | | [10] |
| CENPK+pMW042 | CEN.PK2-1C, 2μ-(pTEF1_FUSn-meGFP-FTH1_tACT1) | FigS1B | This Study |
| CENPK+pMW091, pMW136 | CEN.PK2-1C, 2μ-(pTDH3_SH3D-FUSn-meGFP-FTH1_tADH1,pPGK1_SH3P-miRFP670_tACT1) | FigS2A | This Study |
| CENPK+pMW092, pMW140 | CEN.PK2-1C, 2μ-(pTDH3_PDZD-FUSn-meGFP-FTH1_tADH1,pPGK1_PDZP-miRFP670_tACT1) | FigS2A | This Study |
| yKX013 | CEN.PK2-1C, his3::HIS3Cg-(pTEF1_PDZD-FUSn-mCherry-FTH1_tTDH1), leu2::LEU2Cg-(pHHF2_meGFP-PDZP_tACT1) | Fig1D, Fig1E, Fig2B, Fig4A, FigS2B, FigS3, FigS4 | This Study |
| yKX014 | CEN.PK2-1C, his3::HIS3Cg-(pTEF1_ALFAD-FUSn-mCherry-FTH1_tTDH1), leu2::LEU2Cg-(pHHF2_ALFAP-meGFP_tACT1) | Fig1D, Fig4A, FigS2B, FigS3, FigS4 | This Study |
| yKX015 | CEN.PK2-1C, his3::HIS3Cg-(pTEF1_ALFAD-FUSn-mCherry-FTH1_tTDH1), leu2::LEU2Cg-(pHHF2_meGFP-ALFAP_tACT1) | Fig1D, Fig1E, Fig2B, Fig4A, FigS2B, FigS3, FigS4 | This Study |
| yKX021 | CEN.PK2-1C, his3::HIS3Cg-(pTEF1_SH3D-FUSn-mCherry-FTH1_tTDH1), leu2::LEU2Cg-(pHHF2_meGFP-SH3P_tACT1) | Fig1D, Fig1E, Fig2B, Fig4A, FigS2B, FigS3, FigS4 | This Study |
| yKX024 | CEN.PK2-1C, his3::HIS3Cg-(pTEF1_PTBD-FUSn-mCherry-FTH1_tTDH1), leu2::LEU2Cg-(pHHF2_PTBP-meGFP_tACT1) | Fig1D, Fig4A, FigS2B, FigS3, FigS4 | This Study |
| yKX025 | CEN.PK2-1C, his3::HIS3Cg-(pTEF1_PTBD-FUSn-mCherry-FTH1_tTDH1), leu2::LEU2Cg-(pHHF2_meGFP-PTBP_tACT1) | Fig1D, Fig1E, Fig2B, Fig4A, FigS2B, FigS3, FigS4 | This Study |
| yKX026 | CEN.PK2-1C, his3::HIS3Cg-(pTEF1_scFvD-FUSn-mCherry-FTH1_tTDH1), leu2::LEU2Cg-(pHHF2_scFvP-meGFP_tACT1) | Fig1D, Fig4A, FigS2B, FigS3, FigS4 | This Study |
| yKX027 | CEN.PK2-1C, his3::HIS3Cg-(pTEF1_scFvD-FUSn-mCherry-FTH1_tTDH1), leu2::LEU2Cg-(pHHF2_meGFP-scFvP_tACT1) | Fig1D, Fig1E, Fig2B, Fig4A, FigS2B, FigS3, FigS4 | This Study |
| yKX036 | CEN.PK2-1C, his3::HIS3Cg-(pTEF1_SH3D-FUSn-mCherry-FTH1_tTDH1), leu2::LEU2Cg-(pHHF2_meGFP-PDZP_tACT1) | Fig4A | This Study |
| yKX037 | CEN.PK2-1C, his3::HIS3Cg-(pTEF1_PDZD-FUSn-mCherry-FTH1_tTDH1), leu2::LEU2Cg-(pHHF2_meGFP_tACT1) | Fig4A | This Study |
| yKX038 | CEN.PK2-1C, his3::HIS3Cg-(pTEF1_PDZD-FUSn-mCherry-FTH1_tTDH1), leu2::LEU2Cg-(pHHF2_meGFP-SH3P_tACT1) | Fig4A | This Study |
| yKX039 | CEN.PK2-1C, his3::HIS3Cg-(pTEF1_PDZD-FUSn-mCherry-FTH1_tTDH1), leu2::LEU2Cg-(pHHF2_ALFAP-meGFP_tACT1) | Fig4A | This Study |
| yKX041 | CEN.PK2-1C, his3::HIS3Cg-(pTEF1_PDZD-FUSn-mCherry-FTH1_tTDH1), leu2::LEU2Cg-(pHHF2_PTBP-meGFP_tACT1) | Fig4A | This Study |

| | | | |
|---|---|---|---|
| yKX042 | CEN.PK2-1C, his3::HIS3Cg-(pTEF1_PDZD-FUSn-mCherry-FTH1_tTDH1), leu2::LEU2Cg-(pHHF2_meGFP-PTBP_tACT1) | Fig4A | This Study |
| yKX043 | CEN.PK2-1C, his3::HIS3Cg-(pTEF1_PDZD-FUSn-mCherry-FTH1_tTDH1), leu2::LEU2Cg-(pHHF2_scFvP-meGFP_tACT1) | Fig4A | This Study |
| yKX044 | CEN.PK2-1C, his3::HIS3Cg-(pTEF1_PDZD-FUSn-mCherry-FTH1_tTDH1), leu2::LEU2Cg-(pHHF2_meGFP-scFvP_tACT1) | Fig4A | This Study |
| yKX047 | CEN.PK2-1C, his3::HIS3Cg-(pTEF1_ALFAD-FUSn-mCherry-FTH1_tTDH1), leu2::LEU2Cg-(pHHF2_meGFP-PDZP_tACT1) | Fig4A | This Study |
| yKX049 | CEN.PK2-1C, his3::HIS3Cg-(pTEF1_ALFAD-FUSn-mCherry-FTH1_tTDH1), leu2::LEU2Cg-(pHHF2_meGFP-SH3P_tACT1) | Fig4A | This Study |
| yKX050 | CEN.PK2-1C, his3::HIS3Cg-(pTEF1_ALFAD-FUSn-mCherry-FTH1_tTDH1), leu2::LEU2Cg-(pHHF2_PTBP-meGFP_tACT1) | Fig4A | This Study |
| yKX051 | CEN.PK2-1C, his3::HIS3Cg-(pTEF1_ALFAD-FUSn-mCherry-FTH1_tTDH1), leu2::LEU2Cg-(pHHF2_meGFP-PTBP_tACT1) | Fig4A | This Study |
| yKX052 | CEN.PK2-1C, his3::HIS3Cg-(pTEF1_ALFAD-FUSn-mCherry-FTH1_tTDH1), leu2::LEU2Cg-(pHHF2_scFvP-meGFP_tACT1) | Fig4A | This Study |
| yKX053 | CEN.PK2-1C, his3::HIS3Cg-(pTEF1_ALFAD-FUSn-mCherry-FTH1_tTDH1), leu2::LEU2Cg-(pHHF2_meGFP-scFvP_tACT1) | Fig4A | This Study |
| yKX065 | CEN.PK2-1C, his3::HIS3Cg-(pTEF1_SH3D-FUSn-mCherry-FTH1_tTDH1), leu2::LEU2Cg-(pHHF2_ALFAP-meGFP_tACT1) | Fig4A | This Study |
| yKX066 | CEN.PK2-1C, his3::HIS3Cg-(pTEF1_SH3D-FUSn-mCherry-FTH1_tTDH1), leu2::LEU2Cg-(pHHF2_meGFP-ALFAP_tACT1) | Fig4A | This Study |
| yKX068 | CEN.PK2-1C, his3::HIS3Cg-(pTEF1_SH3D-FUSn-mCherry-FTH1_tTDH1), leu2::LEU2Cg-(pHHF2_PTBP-meGFP_tACT1) | Fig4A | This Study |
| yKX069 | CEN.PK2-1C, his3::HIS3Cg-(pTEF1_SH3D-FUSn-mCherry-FTH1_tTDH1), leu2::LEU2Cg-(pHHF2_meGFP-PTBP_tACT1) | Fig4A | This Study |
| yKX070 | CEN.PK2-1C, his3::HIS3Cg-(pTEF1_SH3D-FUSn-mCherry-FTH1_tTDH1), leu2::LEU2Cg-(pHHF2_scFvP-meGFP_tACT1) | Fig4A | This Study |
| yKX071 | CEN.PK2-1C, his3::HIS3Cg-(pTEF1_SH3D-FUSn-mCherry-FTH1_tTDH1), leu2::LEU2Cg-(pHHF2_meGFP-scFvP_tACT1) | Fig4A | This Study |
| yKX074 | CEN.PK2-1C, his3::HIS3Cg-(pTEF1_PTBD-FUSn-mCherry-FTH1_tTDH1), leu2::LEU2Cg-(pHHF2_meGFP-PDZP_tACT1) | Fig4A | This Study |
| yKX075 | CEN.PK2-1C, his3::HIS3Cg-(pTEF1_PTBD-FUSn-mCherry-FTH1_tTDH1), leu2::LEU2Cg-(pHHF2_ALFAP-meGFP_tACT1) | Fig4A | This Study |
| yKX076 | CEN.PK2-1C, his3::HIS3Cg-(pTEF1_PTBD-FUSn-mCherry-FTH1_tTDH1), leu2::LEU2Cg-(pHHF2_meGFP-ALFAP_tACT1) | Fig4A | This Study |

| | | | |
|---|---|---|---|
| yKX078 | CEN.PK2-1C, his3::HIS3Cg-(pTEF1_PTBD-FUSn-mCherry-FTH1_tTDH1), leu2::LEU2Cg-(pHHF2_meGFP-SH3P_tACT1) | Fig4A | This Study |
| yKX079 | CEN.PK2-1C, his3::HIS3Cg-(pTEF1_PTBD-FUSn-mCherry-FTH1_tTDH1), leu2::LEU2Cg-(pHHF2_scFvP-meGFP_tACT1) | Fig4A | This Study |
| yKX080 | CEN.PK2-1C, his3::HIS3Cg-(pTEF1_PTBD-FUSn-mCherry-FTH1_tTDH1), leu2::LEU2Cg-(pHHF2_meGFP-scFvP_tACT1) | Fig4A | This Study |
| yKX083 | CEN.PK2-1C, his3::HIS3Cg-(pTEF1_scFvD-FUSn-mCherry-FTH1_tTDH1), leu2::LEU2Cg-(pHHF2_meGFP-PDZP_tACT1) | Fig4A | This Study |
| yKX084 | CEN.PK2-1C, his3::HIS3Cg-(pTEF1_scFvD-FUSn-mCherry-FTH1_tTDH1), leu2::LEU2Cg-(pHHF2_ALFAP-meGFP_tACT1) | Fig4A | This Study |
| yKX085 | CEN.PK2-1C, his3::HIS3Cg-(pTEF1_scFvD-FUSn-mCherry-FTH1_tTDH1), leu2::LEU2Cg-(pHHF2_meGFP-ALFAP_tACT1) | Fig4A | This Study |
| yKX087 | CEN.PK2-1C, his3::HIS3Cg-(pTEF1_scFvD-FUSn-mCherry-FTH1_tTDH1), leu2::LEU2Cg-(pHHF2_meGFP-SH3P_tACT1) | Fig4A | This Study |
| yKX088 | CEN.PK2-1C, his3::HIS3Cg-(pTEF1_scFvD-FUSn-mCherry-FTH1_tTDH1), leu2::LEU2Cg-(pHHF2_PTBP-meGFP_tACT1) | Fig4A | This Study |
| yKX089 | CEN.PK2-1C, his3::HIS3Cg-(pTEF1_scFvD-FUSn-mCherry-FTH1_tTDH1), leu2::LEU2Cg-(pHHF2_meGFP-PTBP_tACT1) | Fig4A | This Study |
| yKX114 | CEN.PK2-1C, his3::HIS3Cg-(pTEF1_FUSn-mCherry-FTH1_tTDH1) | Fig1B, Fig1C, FigS1A | This Study |
| yKX115 | CEN.PK2-1C, his3::HIS3Cg-(pTEF1_FUSn-mCherry-FTH1_tTDH1), leu2::LEU2Cg-(pHHF2_meGFP-PDZP_tACT1) | Fig1D, Fig1E, Fig2B, FigS3, FigS4 | This Study |
| yKX116 | CEN.PK2-1C, his3::HIS3Cg-(pTEF1_FUSn-mCherry-FTH1_tTDH1), leu2::LEU2Cg-(pHHF2_ALFAP-meGFP_tACT1) | Fig1E, FigS3, FigS4 | This Study |
| yKX117 | CEN.PK2-1C, his3::HIS3Cg-(pTEF1_FUSn-mCherry-FTH1_tTDH1), leu2::LEU2Cg-(pHHF2_meGFP-ALFAP_tACT1) | Fig1D, Fig1E, Fig2B, FigS3, FigS4 | This Study |
| yKX119 | CEN.PK2-1C, his3::HIS3Cg-(pTEF1_FUSn-mCherry-FTH1_tTDH1), leu2::LEU2Cg-(pHHF2_meGFP-SH3P_tACT1) | Fig1D, Fig1E, Fig2B, FigS3, FigS4 | This Study |
| yKX120 | CEN.PK2-1C, his3::HIS3Cg-(pTEF1_FUSn-mCherry-FTH1_tTDH1), leu2::LEU2Cg-(pHHF2_PTBP-meGFP_tACT1) | Fig1E, FigS3, FigS4 | This Study |
| yKX121 | CEN.PK2-1C, his3::HIS3Cg-(pTEF1_FUSn-mCherry-FTH1_tTDH1), leu2::LEU2Cg-(pHHF2_meGFP-PTBP_tACT1) | Fig1D, Fig1E, Fig2B, FigS3, FigS4 | This Study |
| yKX122 | CEN.PK2-1C, his3::HIS3Cg-(pTEF1_FUSn-mCherry-FTH1_tTDH1), leu2::LEU2Cg-(pHHF2_scFvP-meGFP_tACT1) | Fig1E, FigS3, FigS4 | This Study |
| yKX123 | CEN.PK2-1C, his3::HIS3Cg-(pTEF1_FUSn-mCherry-FTH1_tTDH1), leu2::LEU2Cg-(pHHF2_meGFP-scFvP_tACT1) | Fig1D, Fig1E, Fig2B, FigS3, FigS4 | This Study |
| yKX156 | CEN.PK2-1C, his3::HIS3Cg-(pTEF1_ALFAD-FUSn-mCherry-FTH1_tTDH1), leu2::LEU2Cg-(pHHF2_meGFP_tACT1) | Fig4A | This Study |

| | | | |
|---|---|---|---|
| yKX157 | CEN.PK2-1C, his3::HIS3Cg-(pTEF1_SH3D-FUSn-mCherry-FTH1_tTDH1), leu2::LEU2Cg-(pHHF2_meGFP_tACT1) | Fig4A | This Study |
| yKX158 | CEN.PK2-1C, his3::HIS3Cg-(pTEF1_PTBD-FUSn-mCherry-FTH1_tTDH1), leu2::LEU2Cg-(pHHF2_meGFP_tACT1) | Fig4A | This Study |
| yKX159 | CEN.PK2-1C, his3::HIS3Cg-(pTEF1_scFvD-FUSn-mCherry-FTH1_tTDH1), leu2::LEU2Cg-(pHHF2_meGFP_tACT1) | Fig4A | This Study |
| yKX485 | CEN.PK2-1C, YARCdelta5-(pTDH3-EfMvaE-tADH1, pTEF1-EfMvaS-tACT1, pPGK1-SeAcs(L641P)-tCYC1), ΔERG20-(ERG20-Myc-PDZP) | | This Study |
| yKX604 | CEN.PK2-1C, YARCdelta5-(pTDH3-EfMvaE-tADH1, pTEF1-EfMvaS-tACT1, pPGK1-SeAcs(L641P)-tCYC1), ΔERG20-(ERG20-Myc-PDZP), his3::HIS3Cg-(pTDH3_PDZD-PDZD-FUSn-mCherry-I301(K129A)_tTDH1, pTEF1_PDZD-PDZD-FUSn-mCherry-I301(K129A)_tTDH1), leu2::LEU2Cg-(pTEF1_aaFSmut-Myc-SH3P_tSSA1, pTDH3_SH3D-SH3D-FUSn-mCherry-I301(K129A)_tTDH1, pTDH3_aaFSmut-Myc-SH3P_tACT1, pCCW12_SH3D-SH3D-FUSn-mCherry-I301(K129A)_tENO1) | | This Study |
| yKX617 | CEN.PK2-1C, YARCdelta5-(pTDH3-EfMvaE-tADH1, pTEF1-EfMvaS-tACT1, pPGK1-SeAcs(L641P)-tCYC1), ΔERG20-(ERG20-Myc-PDZP), his3::HIS3Cg-(pTDH3_FUSn-mCherry-I301(K129A)_tADH1t, pTEF1p_FUSn-mCherry-I301(K129A)_tTDH1 ), leu2::LEU2Cg-(pTEF1_aaFSmut-Myc-SH3P_tSSA1, pTDH3_FUSn-mCherry-I301(K129A)_tTDH1, pTDH3_aaFSmut-Myc-SH3P_tACT1, pCCW12_FUSn-mCherry-I301(K129A)_tENO1) | Fig4E | This Study |
| yKX623 | CEN.PK2-1C, YARCdelta5-(pTDH3-EfMvaE-tADH1, pTEF1-EfMvaS-tACT1, pPGK1-SeAcs(L641P)-tCYC1), ΔERG20-(ERG20-Myc-PDZP), his3::HIS3Cg-(pTDH3_PDZD-PDZD-FUSn-mCherry-I301(K129A)_tTDH1, pTEF1_PDZD-PDZD-FUSn-mCherry-I301(K129A)_tTDH1) | | This Study |
| yKX624 | CEN.PK2-1C, YARCdelta5-(pTDH3-EfMvaE-tADH1, pTEF1-EfMvaS-tACT1, pPGK1-SeAcs(L641P)-tCYC1), ΔERG20-(ERG20-Myc-PDZP), his3::HIS3Cg-(pTDH3_FUSn-mCherry-I301(K129A)_tADH1t, pTEF1p_FUSn-mCherry-I301(K129A)_tTDH1 ) | | This Study |

| | | | |
|---|---|---|---|
| yKX646 | CEN.PK2-1C, YARCdelta5-(pTDH3-EfMvaE-tADH1, pTEF1-EfMvaS-tACT1, pPGK1-SeAcs(L641P)-tCYC1), ΔERG20-(ERG20-Myc-PDZP), his3::HIS3Cg-(pTDH3_PDZD-PDZD-FUSn-mCherry-I301(K129A)_tTDH1, pTEF1_PDZD-PDZD-FUSn-mCherry-I301(K129A)_tTDH1), leu2::LEU2Cg-(pTEF1_aaFSmut-Myc-SH3P_tSSA1, pTDH3_SH3D-SH3D-FUSn-mCherry-I301(K129A)_tTDH1, pTDH3_aaFSmut-Myc-SH3P_tACT1, pCCW12_SH3D-SH3D-FUSn-mCherry-I301(K129A)_tENO1), XII-2::KanMX-(pTDH3_SH3D-SH3D-FUSn-mCherry-I301(K129A)_tTDH1, pCCW12_SH3D-SH3D-FUSn-mCherry-I301(K129A)_tENO1) | Fig4E | This Study |
| yKX647 | CEN.PK2-1C, YARCdelta5-(pTDH3-EfMvaE-tADH1, pTEF1-EfMvaS-tACT1, pPGK1-SeAcs(L641P)-tCYC1), ΔERG20-(ERG20-Myc-PDZP), his3::HIS3Cg-(pTDH3_PDZD-PDZD-FUSn-mCherry-I301(K129A)_tTDH1, pTEF1_PDZD-PDZD-FUSn-mCherry-I301(K129A)_tTDH1), leu2::LEU2Cg-(pTEF1_aaFSmut-Myc-SH3P_tSSA1, pTDH3_SH3D-SH3D-FUSn-mCherry-I301(K129A)_tTDH1, pTDH3_aaFSmut-Myc-SH3P_tACT1, pCCW12_SH3D-SH3D-FUSn-mCherry-I301(K129A)_tENO1), XII-2::KanMX-(pCCW12_PDZD-PDZD-FUSn-mCherry-I301(K129A)_tENO1 pTDH3_PDZD-PDZD-FUSn-mCherry-I301(K129A)_tTDH1) | Fig4E | This Study |
| yKX656 | CEN.PK2-1C, YARCdelta5-(pTDH3-EfMvaE-tADH1, pTEF1-EfMvaS-tACT1, pPGK1-SeAcs(L641P)-tCYC1), ΔERG20-(ERG20-Myc-PDZP), his3::HIS3Cg-(pTDH3_PDZD-PDZD-FUSn-mCherry-I301(K129A)_tTDH1, pTEF1_PDZD-PDZD-FUSn-mCherry-I301(K129A)_tTDH1), leu2::LEU2Cg-(pTEF1_aaFSmut-Myc-SH3P_tSSA1, pTDH3_SH3D-SH3D-FUSn-mCherry-I301(K129A)_tTDH1, pTDH3_aaFSmut-Myc-SH3P_tACT1, pCCW12_SH3D-SH3D-FUSn-mCherry-I301(K129A)_tENO1), XII-2::KanMX-(pCCW12_PDZD-PDZD-FUSn-mCherry-I301(K129A)_ENO1t pTDH3_SH3D-SH3D-FUSn-mCherry-I301(K129A)_tTDH1) | Fig4E | This Study |
| yMW703 | CEN.PK2-1C, his3::HIS3Cg-(pTEF1_FUSn-mCherry-FTH1_tTDH1), leu2::LEU2Cg-(pTEF1_FUSn-mCherry-FTH1_tTDH1) | Fig1B, Fig1C, FigS1A | This Study |
| yMW766 | CEN.PK2-1C, his3::HIS3Cg-(pTEF1_FUSn-mCherry-FTH1_tTDH1), leu2::LEU2Cg-(pTEF1_FUSn-mCherry-FTH1_tTDH1), trp1::TRP1Cg-(pTEF1_FUSn-mCherry-FTH1_tTDH1) | Fig1B, Fig1C, FigS1A | This Study |

| | | | |
|---|---|---|---|
| yMW778 | CEN.PK2-1C, his3::HIS3Cg-(pTEF1_FUSn-mCherry-FTH1_tTDH1), leu2::LEU2Cg-(pTEF1_FUSn-mCherry-FTH1_tTDH1), trp1::TRP1Cg-(pTEF1_FUSn-mCherry-FTH1_tTDH1), CEN-(pTEF1_FUSn-mCherry-FTH1_tACT1) | Fig1B, Fig1C, FigS1A | This Study |
| yMW878 | CEN.PK2-1C, leu2::LEU2Cg-(pRPL18B_meGFP-PTBP_tACT1), his3::HIS3Cg-(pTDH3_PTBD-FUSn-mCherry-FTH1_tTDH1) | Fig3FG(ii), FigS6, FigS7 | This Study |
| yMW879 | CEN.PK2-1C, leu2::LEU2Cg-(pHHF2_meGFP-PTBP_tACT1), his3::HIS3Cg-(pTDH3_PTBD-FUSn-mCherry-FTH1_tTDH1) | Fig3FG(ii), FigS7 | This Study |
| yMW880 | CEN.PK2-1C, leu2::LEU2Cg-(pTEF1_meGFP-PTBP_tACT1), his3::HIS3Cg-(pTDH3_PTBD-FUSn-mCherry-FTH1_tTDH1) | Fig3FG(ii), FigS7 | This Study |
| yMW884 | CEN.PK2-1C, leu2::LEU2Cg-(pRPL18B_meGFP-PTBP_tACT1), his3::HIS3Cg-(pHHF2_PTBD-FUSn-mCherry-FTH1_tTDH1) | Fig3FG(ii), FigS7 | This Study |
| yMW887 | CEN.PK2-1C, leu2::LEU2Cg-(pHHF2_meGFP-PTBP_tACT1), his3::HIS3Cg-(pHHF2_PTBD-FUSn-mCherry-FTH1_tTDH1) | Fig3FG(ii), FigS7 | This Study |
| yMW890 | CEN.PK2-1C, leu2::LEU2Cg-(pTEF1_meGFP-PTBP_tACT1), his3::HIS3Cg-(pHHF2_PTBD-FUSn-mCherry-FTH1_tTDH1) | Fig3FG(ii), FigS7 | This Study |
| yMW918 | CEN.PK2-1C, leu2::LEU2Cg-(pRPL18B_meGFP-PTBP_tACT1), his3::HIS3Cg-(pTDH3_PTBD-FUSn-mCherry-FTH1_tENO1, pTDH3_PTBD-FUSn-mCherry-FTH1_tTDH1) | Fig3FG(ii), FigS7 | This Study |
| yMW919 | CEN.PK2-1C, leu2::LEU2Cg-(pHHF2_meGFP-PTBP_tACT1), his3::HIS3Cg-(pTDH3_PTBD-FUSn-mCherry-FTH1_tENO1, pTDH3_PTBD-FUSn-mCherry-FTH1_tTDH1) | Fig3FG(ii), FigS7 | This Study |
| yMW920 | CEN.PK2-1C, leu2::LEU2Cg-(pTEF1_meGFP-PTBP_tACT1), his3::HIS3Cg-(pTDH3_PTBD-FUSn-mCherry-FTH1_tENO1, pTDH3_PTBD-FUSn-mCherry-FTH1_tTDH1) | Fig3FG(ii), FigS7 | This Study |
| yMW966 | CEN.PK2-1C, leu2::LEU2Cg-(pHHF2_ALFAP-Myc-VenusC_tACT1, pHHF2_VenusN-HA-PTBP_tADH1), his3::HIS3Cg-(pCCW12_ALFAD-FUSn-mCherry-FTH1_tENO1, pHHF2_PTBD-FUSn-mCherry-FTH1_tTDH1) | Fig4C | This Study |
| yMW983 | CEN.PK2-1C, leu2::LEU2Cg-(pHHF2_meGFP-PDZP_tACT1), his3::HIS3Cg-(pTDH3_PDZD-FUSn-mCherry-FTH1_tTDH1) | FigS5 | This Study |
| yMW987 | CEN.PK2-1C, leu2::LEU2Cg-(pHHF2_meGFP-PDZP_tACT1), his3::HIS3Cg-(pTDH3_PDZD-FUSn(3YS)-mCherry-FTH1_tTDH1) | FigS5 | This Study |
| yMW988 | CEN.PK2-1C, leu2::LEU2Cg-(pHHF2_meGFP-PDZP_tACT1), his3::HIS3Cg-(pTDH3_PDZD-FUSn(5YS)-mCherry-FTH1_tTDH1) | FigS5 | This Study |
| yMW989 | CEN.PK2-1C, leu2::LEU2Cg-(pHHF2_meGFP-PDZP_tACT1), his3::HIS3Cg-(pTDH3_PDZD-FUSn(9YS)-mCherry-FTH1_tTDH1) | FigS5 | This Study |

| | | | |
|---|---|---|---|
| yMW990 | CEN.PK2-1C, leu2::LEU2Cg-(pHHF2_meGFP-PDZP_tACT1), his3::HIS3Cg-(pTDH3_PDZD-FUSn(15YS)-mCherry-FTH1_tTDH1) | FigS5 | This Study |
| yMW991 | CEN.PK2-1C, leu2::LEU2Cg-(pHHF2_meGFP-PDZP_tACT1), his3::HIS3Cg-(pTDH3_PDZD-FUSn(27YS)-mCherry-FTH1_tTDH1) | FigS5 | This Study |
| yMW1026 | CEN.PK2-1C, leu2::LEU2Cg-(pHHF2_ALFAP-Myc-VenusC_tACT1, pHHF2_VenusN-HA-PTBP_tADH1), his3::HIS3Cg-(pTDH3_FUSn-mCherry-FTH1_tTDH1, pCCW12_FUSn-mCherry-FTH1_tENO1) | Fig4C | This Study |
| yMW1027 | CEN.PK2-1C, leu2::LEU2Cg-(pHHF2_ALFAP-Myc-VenusC_tACT1, pHHF2_VenusN-HA-PTBP_tADH1), his3::HIS3Cg-(pCCW12_ALFAD-FUSn-mCherry-FTH1_tENO1, pTDH3_FUSn-mCherry-FTH1_tTDH1t ) | Fig4C | This Study |
| yMW1028 | CEN.PK2-1C, leu2::LEU2Cg-(pHHF2_ALFAP-Myc-VenusC_tACT1, pHHF2_VenusN-HA-PTBP_tADH1), his3::HIS3Cg-(pCCW12_FUSn-mCherry-FTH1_tENO1, pTDH3_PTBD-FUSn-mCherry-FTH1_tTDH1) | Fig4C | This Study |
| yMW1038 | CEN.PK2-1C, leu2::LEU2Cg-(pHHF2_ALFAP-Myc-VenusC_tACT1, pHHF2_VenusN-HA-PTBP_tADH1), his3::HIS3Cg-(pCCW12_ALFAD-FUSn-mCherry-FTH1_tENO1, pTDH3_PTBD-FUSn-mCherry-FTH1_tTDH1) | Fig4C | This Study |
| yMW1048 | CEN.PK2-1C, leu2::LEU2Cg-(pHHF2_ALFAP-Myc-VenusC_tACT1, pHHF2_VenusN-HA-PTBP_tADH1), his3::HIS3Cg-() | Fig4C | This Study |
| yMW1143 | CEN.PK2-1C, his3::HIS3Cg-(pTDH3_PTBD-FUSn-mCherry-FTH1_tTDH1), leu2::LEU2Cg-(pHHF2_BCRDimer-meGFP-PTBP_tACT1) | FigS6 | This Study |
| yMW1146 | CEN.PK2-1C, his3::HIS3Cg-(pTDH3_PTBD-FUSn-mCherry-FTH1_tTDH1), leu2::LEU2Cg-(pHHF2_CCTrimer-meGFP-PTBP_tACT1) | FigS6 | This Study |
| yMW1150 | CEN.PK2-1C, his3::HIS3Cg-(pTDH3_PTBD-FUSn-mCherry-FTH1_tTDH1), leu2::LEU2Cg-(pRPL18B_meGFP-PTBP-PTBP_tACT1) | Fig3FG(i), FigS7 | This Study |
| yMW1151 | CEN.PK2-1C, his3::HIS3Cg-(pTDH3_PTBD-FUSn-mCherry-FTH1_tTDH1), leu2::LEU2Cg-(pHHF2_meGFP-PTBP-PTBP_tACT1) | Fig3FG(i), FigS7 | This Study |
| yMW1152 | CEN.PK2-1C, his3::HIS3Cg-(pTDH3_PTBD-FUSn-mCherry-FTH1_tTDH1), leu2::LEU2Cg-(pTEF1_meGFP-PTBP-PTBP_tACT1) | Fig3FG(i), FigS7 | This Study |
| yMW1182 | CEN.PK2-1C, his3::HIS3Cg-(pHHF2_PTBD-FUSn-mCherry-FTH1_tTDH1), leu2::LEU2Cg-(pRPL18B_meGFP-PTBP-PTBP_tACT1) | Fig3FG(i), FigS7 | This Study |
| yMW1183 | CEN.PK2-1C, his3::HIS3Cg-(pHHF2_PTBD-FUSn-mCherry-FTH1_tTDH1), leu2::LEU2Cg-(pHHF2_meGFP-PTBP-PTBP_tACT1) | Fig3FG(i), FigS7 | This Study |
| yMW1184 | CEN.PK2-1C, his3::HIS3Cg-(pHHF2_PTBD-FUSn-mCherry-FTH1_tTDH1), leu2::LEU2Cg-(pTEF1_meGFP-PTBP-PTBP_tACT1) | Fig3FG(i), FigS7 | This Study |

| | | | |
|---|---|---|---|
| yMW1188 | CEN.PK2-1C, his3::HIS3Cg-(pTDH3_PTBD-FUSn-mCherry-FTH1_tENO1, pTDH3_PTBD-FUSn-mCherry-FTH1_tTDH1), leu2::LEU2Cg-(pRPL18B_meGFP-PTBP-PTBP_tACT1) | Fig3FG(i), FigS7 | This Study |
| yMW1189 | CEN.PK2-1C, his3::HIS3Cg-(pTDH3_PTBD-FUSn-mCherry-FTH1_tENO1, pTDH3_PTBD-FUSn-mCherry-FTH1_tTDH1), leu2::LEU2Cg-(pHHF2_meGFP-PTBP-PTBP_tACT1) | Fig3FG(i), FigS7 | This Study |
| yMW1190 | CEN.PK2-1C, his3::HIS3Cg-(pTDH3_PTBD-FUSn-mCherry-FTH1_tENO1, pTDH3_PTBD-FUSn-mCherry-FTH1_tTDH1), leu2::LEU2Cg-(pTEF1_meGFP-PTBP-PTBP_tACT1) | Fig3FG(i), FigS7 | This Study |
| yMW1199 | CEN.PK2-1C, his3::HIS3Cg-(pTDH3_mCherry-(ERBV-1)2A-GFP_tTDH1) | | This Study |

Supplemental Information References
